# Structural basis for the absence of low-energy chlorophylls responsible for photoprotection from a primitive cyanobacterial PSI

**DOI:** 10.1101/2021.09.29.462462

**Authors:** Koji Kato, Tasuku Hamaguchi, Ryo Nagao, Keisuke Kawakami, Yoshifumi Ueno, Takehiro Suzuki, Hiroko Uchida, Akio Murakami, Yoshiki Nakajima, Makio Yokono, Seiji Akimoto, Naoshi Dohmae, Koji Yonekura, Jian-Ren Shen

**Author notes:** These authors contributed equally to this work (Koji Kato, Tasuku Hamaguchi, Ryo Nagao). Corresponding Authors: Ryo Nagao, Koji Yonekura, Jian-Ren Shen.

## Abstract

Photosystem I (PSI) of photosynthetic organisms is a multi-subunit pigment-protein complex and functions in light harvesting and photochemical charge-separation reactions, followed by reduction of NADP to NADPH required for CO_2_ fixation. PSI from different photosynthetic organisms has a variety of chlorophylls (Chls), some of which are at lower-energy levels than its reaction center P700, a special pair of Chls, and are called low-energy Chls. However, the site of low-energy Chls is still under debate. Here, we solved a 2.04-Å resolution structure of a PSI trimer by cryo-electron microscopy from a primitive cyanobacterium *Gloeobacter violaceus* PCC 7421, which has no low-energy Chls. The structure showed absence of some subunits commonly found in other cyanobacteria, confirming the primitive nature of this cyanobacterium. Comparison with the known structures of PSI from other cyanobacteria and eukaryotic organisms reveals that one dimeric and one trimeric Chls are lacking in the *Gloeobacter* PSI. The dimeric and trimeric Chls are named Low1 and Low2, respectively. Low2 does not exist in some cyanobacterial and eukaryotic PSIs, whereas Low1 is absent only in *Gloeobacter*. Since *Gloeobacter* is susceptible to light, this indicates that Low1 serves as a main photoprotection site in most oxyphototrophs, whereas Low2 is involved in either energy transfer or energy quenching in some of the oxyphototrophs. Thus, these findings provide insights into not only the functional significance of low-energy Chls in PSI, but also the evolutionary changes of low-energy Chls responsible for the photoprotection machinery from photosynthetic prokaryotes to eukaryotes.

## Introduction

Oxygenic photosynthetic reactions convert light energy into chemical energy and produce molecular oxygen, thereby maintaining the aerobic life on the earth (1). Despite the necessity of light energy in photosynthesis, excess-light energy is harmful to the photosynthetic reactions (2–4). Therefore, oxyphototrophs have developed various photoprotection mechanisms to deal with excess-light energy, and photoprotection is essential for oxyphototrophs to survive under changeable light conditions. One of the photoprotection mechanisms is excitation-energy quenching by converting excitation energy to harmless heat (5–7). Energy quenching occurs mainly by either energy transfer or charge transfer between chlorophylls (Chls) and carotenoids (Cars) (4, 8–10).

The light-induced photosynthetic reactions take place in two multi-subunit pigment-protein complexes, photosystem I and photosystem II (PSI and PSII, respectively). PSI is a multi-subunit pigment-protein complex and functions in light harvesting, charge-separation and electron transfer reactions, and in reduction of NADP to NADPH required for CO_2_ fixation (1, 11, 12). PSI has many cofactors including chlorophylls (Chls), carotenoids (Cars), quinones, and iron-sulfur clusters functioning in the photochemical reactions (12). These cofactors are well conserved in PSI cores among oxyphototrophs, although the molecular organizations of PSI differ significantly among different species of organisms (13).

Among photosynthetic organisms, cyanobacteria are known to be a large group of prokaryotes and to have either trimeric (14–19) or tetrameric (20–22) PSI complexes. Cyanobacterial PSI complexes also possess unique Chl molecules at energy levels lower than the level of P700, a special pair Chl that performs charge separation among ~95 Chl molecules in the PSI-monomer unit (11). The low-energy Chls are historically called Red Chls, which have different energy levels among cyanobacteria (23, 24). The major functions of the low-energy Chls in PSI are either uphill energy transfer from low-energy Chls to other Chls (23) or excitation-energy quenching under excess excitation conditions (25–27). From these observations, low-energy Chls are thought to regulate energy balance for energy transfer and energy quenching (23, 24). Despite numerous structural and functional analyses of the cyanobacterial PSI, the location of low-energy Chls is still under debate for a long time both experimentally and theoretically (23, 24, 28, 29).

The low-energy Chls in PSI are characterized by their clear spectral components using fluorescence spectroscopy at 77 K. There are two types of major fluorescence peaks from low-energy Chls which are located at around 723 and 730 nm in PSI trimers isolated from the representative cyanobacteria, *Synechocystis* sp. PCC 6803 (hereafter referred to as *Synechocystis*) (30, 31) and *Thermosynechococcus elongatus* (32), respectively. The fluorescence feature of low-energy Chls is conserved even in the tetrameric PSI cores of cyanobacteria (20, 33). In this study, we define the 723-nm and 730-nm low-energy Chls as Low1 and Low2, respectively. Most of cyanobacteria have Low1 and/or Low2; however, it is very interesting to note that the cyanobacterium *Gloeobacter violaceus* PCC 7421 (hereafter referred to as *Gloeobacter*) possesses neither Low1 nor Low2 in PSI as revealed by fluorescence analyses of *in vivo* (34) and *in vitro* (31). *Gloeobacter* has no thylakoid membranes (35) and branches off from the main cyanobacterial tree at an early state (36), thereby is classified as a primitive cyanobacterium. An additional feature of *Gloeobacter* is its susceptibility to light (37, 38). *Gloeobacter* cannot grow under a normal-light condition for other cyanobacteria, e.g., 30 μmol photons m^−2^ s^−1^, under which, most oxyphototrophs including *Synechocystis* and *T. elongatus* grow well. This means that *Gloeobacter* does not possess a functional photoprotection mechanism present in *Synechocystis* and *T. elongatus*. In other words, if this photoprotection mechanism disappears, oxyphototrophs will die under normal-light conditions. These characteristics, namely, the absence of low-energy Chls in PSI and the ability of *Gloeobacter* to grow only under extremely low light conditions lead to a notion that Low1 and/or Low2 are involved in the photoprotection mechanism.

In order to reveal the locations of Low1 and Low2 in *Synechocystis* and thermophilic cyanobacteria, we solved the structure of the PSI trimer isolated from *Gloeobacter* using cryo-electron microscopy (cryo-EM) at a resolution of 2.04 Å. The structure obtained showed the absence of some subunits commonly found in other cyanobacteria, confirming the primitive nature of this cyanobacterium. By comparing the *Gloeobacter* PSI structure with the structures of *Synechocystis* and thermophilic cyanobacteria, we were able to locate the positions of Low1 and Low2 sites; based on which we discuss the correlation of Low1 and Low2 with the photoprotection machinery required for survival of most oxyphototrophs under normal-light conditions.

## Results

### Overall structure of the PSI trimer

The PSI trimers were purified from *Gloeobacter*, and its biochemical characterization was summarized in Fig. S1. Cryo-EM images of the PSI trimer were obtained by a JEOL CRYO ARM 300 electron microscope operated at 300 kV. After processing of the images with RELION (Fig. S2 and Table S1), the final cryo-EM map with a C3 symmetry enforced was determined at a resolution of 2.04 Å, based on the “gold standard” Fourier shell correlation (FSC) = 0.143 criterion (Fig. 1*A* and Fig. S2). This resolution is the highest for the structure of PSI trimers ever determined by X-ray crystallography and single particle cryo-EM so far (14–19). This was realized by imaging with a cold-field emission electron beam that produces superior high-resolution signals (39). The atomic model of PSI was built based on the 2.04-Å map (Fig. 1*B* and Table S2), and most of the cofactors and amino acid residues were precisely assigned into this high-resolution map.

**Fig. 1.**
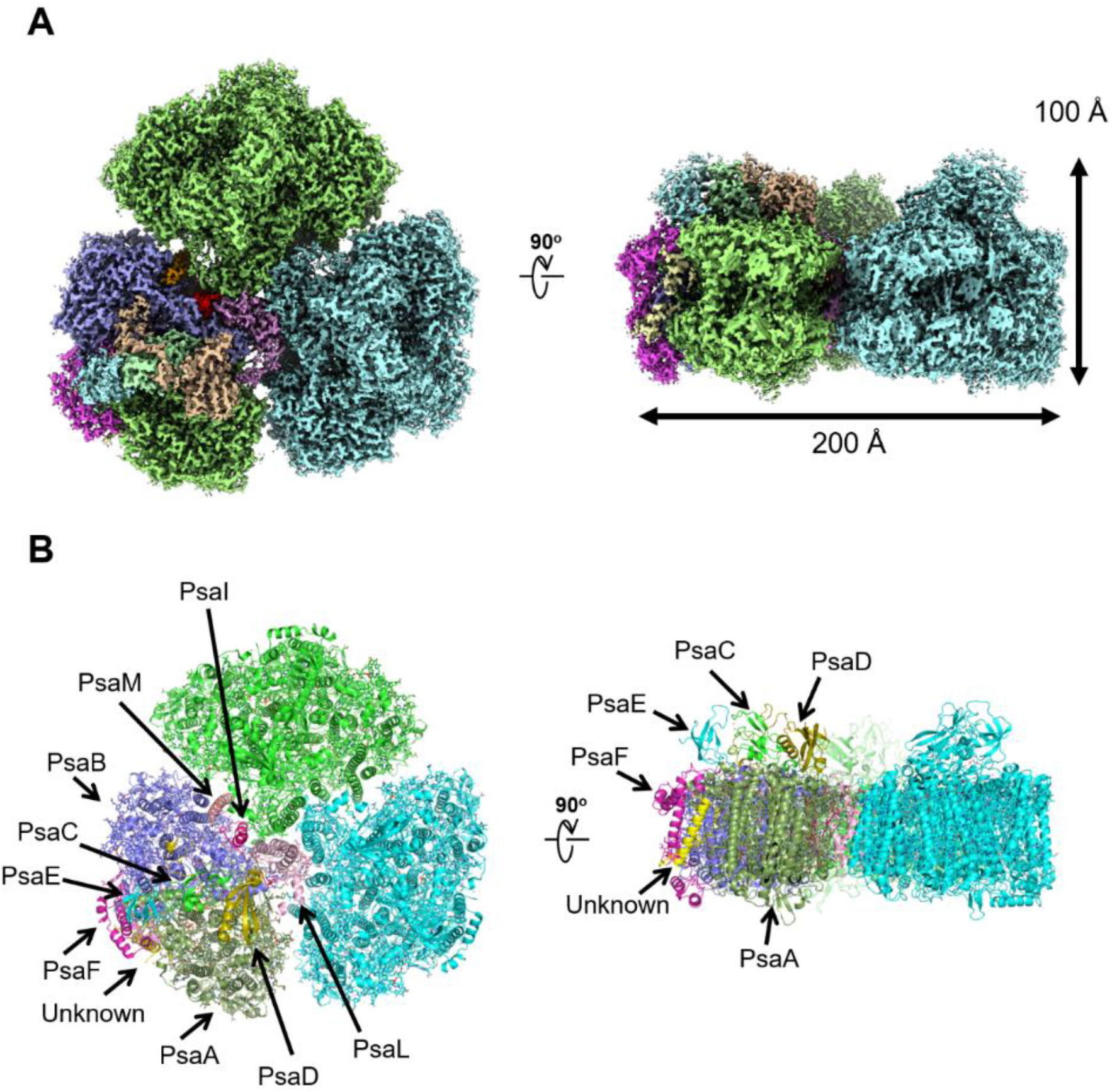
Overall structure of the *Gloeobacter* PSI trimer. (**A**) The cryo-EM density of PSI at 2.04 Å resolution. (**B**) Structural model of the PSI trimer. Views are from the top of the cytosolic side (left) and side of the membrane (right) for both panels **A** and **B**.

The electron-transfer chains of a special pair Chls P700, accessary Chl Acc, primary electron acceptor A0, menaquinone-4 A_1_, and iron-sulfur clusters FX/FA/FB are assigned unambiguously (Fig. 2). The densities for the Mg atoms in the Chl molecules are clearly visualized at their centers (Figs. 2*B*–2*E*). In addition, the characteristic structures between Chl *a*′ and Chl *a* in P700 are distinguished clearly in the high-resolution map (red arrows in Fig 2*B* and 2*C*, respectively). The densities of individual heavy atoms in the iron-sulfur clusters are also clearly separated, allowing their precise assignment possible (Figs. 2*G*–2*I*).

**Fig. 2.**
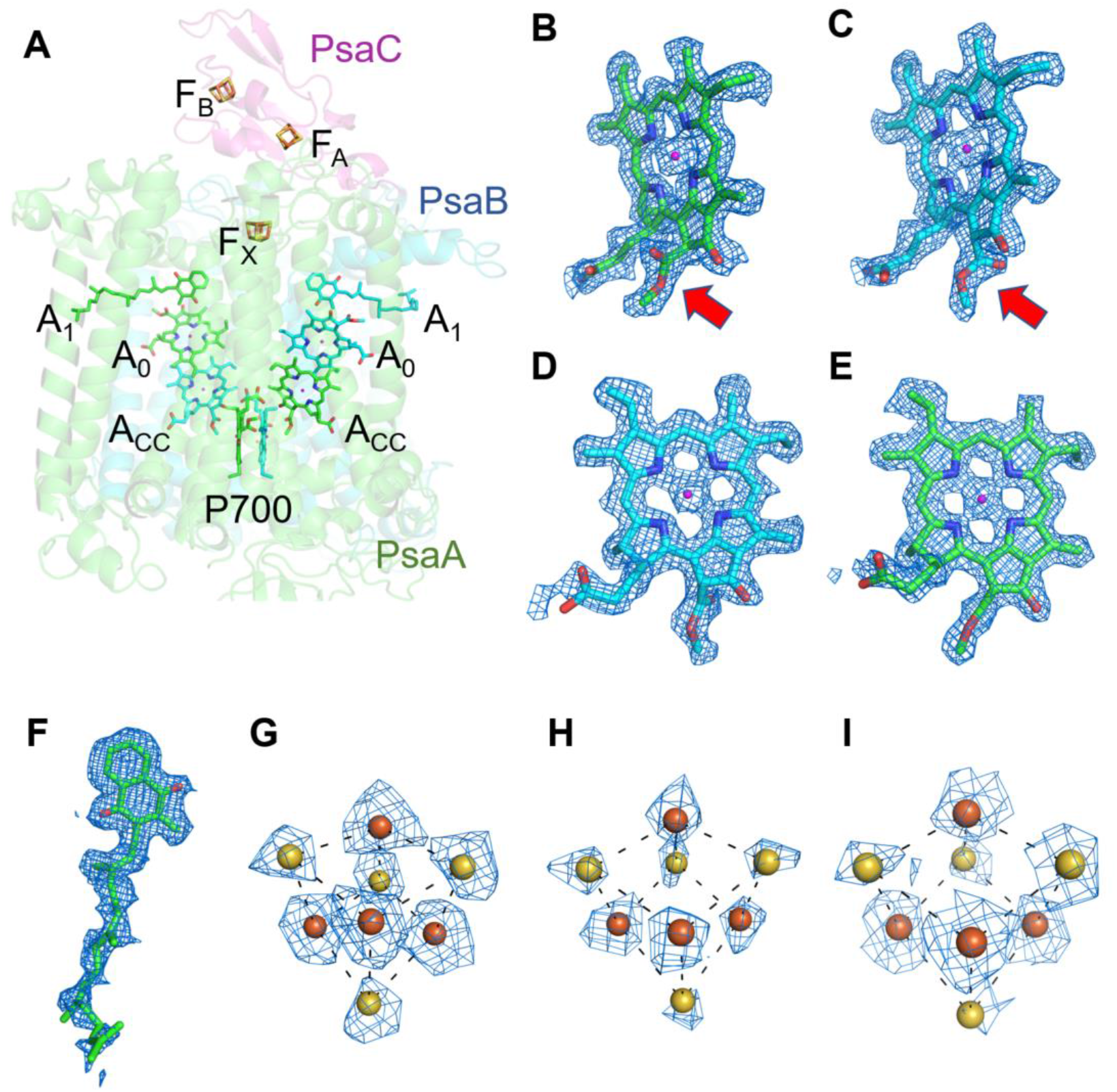
Cofactors involved in electron-transfer reaction in the *Gloeobacter* PSI. (**A**) Arrangement of cofactors involved in the electron-transfer reaction. P700, special pair Chls; Acc, accessary Chl; A_0_, primary electron acceptor; A_1_, menaquinone-4; and F_X_, F_A_, and F_B_, iron-sulfur clusters. (**B**–**I**) The cryo-EM density maps of cofactors and their refined models. (**B**) Chl *a*′ in P700; (**C**) Chl *a* in P700; (**D**) Acc; (**E**) A_0_; (**F**) A_1_; and (**G**–**I**), F_X_, F_A_, and F_B_. Red arrows indicate structural differences between Chl *a* and Chl *a*′. (**B**, **C**). The densities for Chls, quinone, and iron-sulfur clusters were depicted at 5 σ, 3 σ, and 15 σ, respectively.

### Subunit structures of the PSI monomer

PSI of *Gloeobacter* is a homo-trimeric complex, and its overall architecture is similar to those of the PSI trimer isolated from other cyanobacteria (14–19). Each monomer of the *Gloeobacter* PSI contains nine subunits (Fig. 1*B*), and all of the genes for these subunits are found in the genome of *Gloeobacter* (*psaA*, *psaB*, *psaC*, *psaD*, *psaE*, *psaF*, *psaL*, *psaM*, and *psaZ) (40).* PsaZ was positioned at a same location of PsaI in the PSI structure of *Synechocystis* (15), although the *Gloeobacter* PsaZ has low sequence identity (20.0%) with the *Synechocystis* PsaI (Figs. S3*A* and S3*B*) (41). The other eight subunits were located at similar positions of PSI from other cyanobacteria (14–19). An additional subunit was found at the position of PsaJ which was modelled as polyalanines (Fig. S3*C*), because the *psaJ* gene is absent in the *Gloeobacter* genome (40) but a subunit is clearly visible in the map at the same position as PsaJ (14–19). This subunit is named Unknown in the *Gloeobacter* structure. The cryo-EM map of *Gloeobacter* PSI indicated the absence of PsaG, PsaH, PsaK, and PsaX found in other PSIs, consistent with the absence of their genes in the *Gloeobacter* genome (40). While PsaG and PsaH are also absent in other cyanobacteria and appeared in the green lineage PSI, PsaK and PsaX are found in other cyanobacterial PSI and binds Chls (see below) and play important roles in stabilizing the PSI structure and in energy transfer (13–19). The absence of these subunits in the *Gloeobacter* PSI is consistent with its primitive nature during the evolutionary process.

The root mean square deviations between the monomer unit of *Gloeobacter* PSI and that of cyanobacterial PSI with the same set of subunits from *T. elongatus* (PDB: 1JB0) (14), *Synechocystis* (PDB: 5OY0) (15), *T. vulcanus* (PDB: 6K33) (16), *Halomicronema hongdechloris* (PDB: 6KMW) (17), and *Acaryochloris marina* (PDB: 7COY) (18), are 0.85 Å for 2,008 Cα atoms, 0.82 Å for 2,019 Cα atoms, 0.90 Å for 2,011 Cα atoms, 0.88 Å for 1,785 Cα atoms, and 1.10 Å for 1,896 Cα atoms, respectively. This suggests that the overall structures of PSI are largely similar among the cyanobacteria, if we do not include some subunits that are absent or different between different cyanobacteria. However, some regions of the structure of *Gloeobacter* PSI display large differences compared with other cyanobacterial PSI (Fig. 3 and Figs. S4–S6). The *Gloeobacter* PSI possesses four types of characteristic loop structures which are named Loop1 (Tyr515–Gln529) and Loop2 (Asn652– Ser665) in PsaA, Loop3 (Pro717–Ile727) in PsaB, and Loop4 (Gln31–Asp36) in PsaF (Fig. 3*A*). These four types of loops do not exist in other cyanobacterial PSI trimers (14–19). In contrast, the motif of Pro237–Gln248 in the *Synechocystis* PsaB is absent in the *Gloeobacter* PSI (Fig. S5). All these insertions and deletions are located at the periplasmic side of the PSI monomer (Fig. 3).

**Fig. 3.**
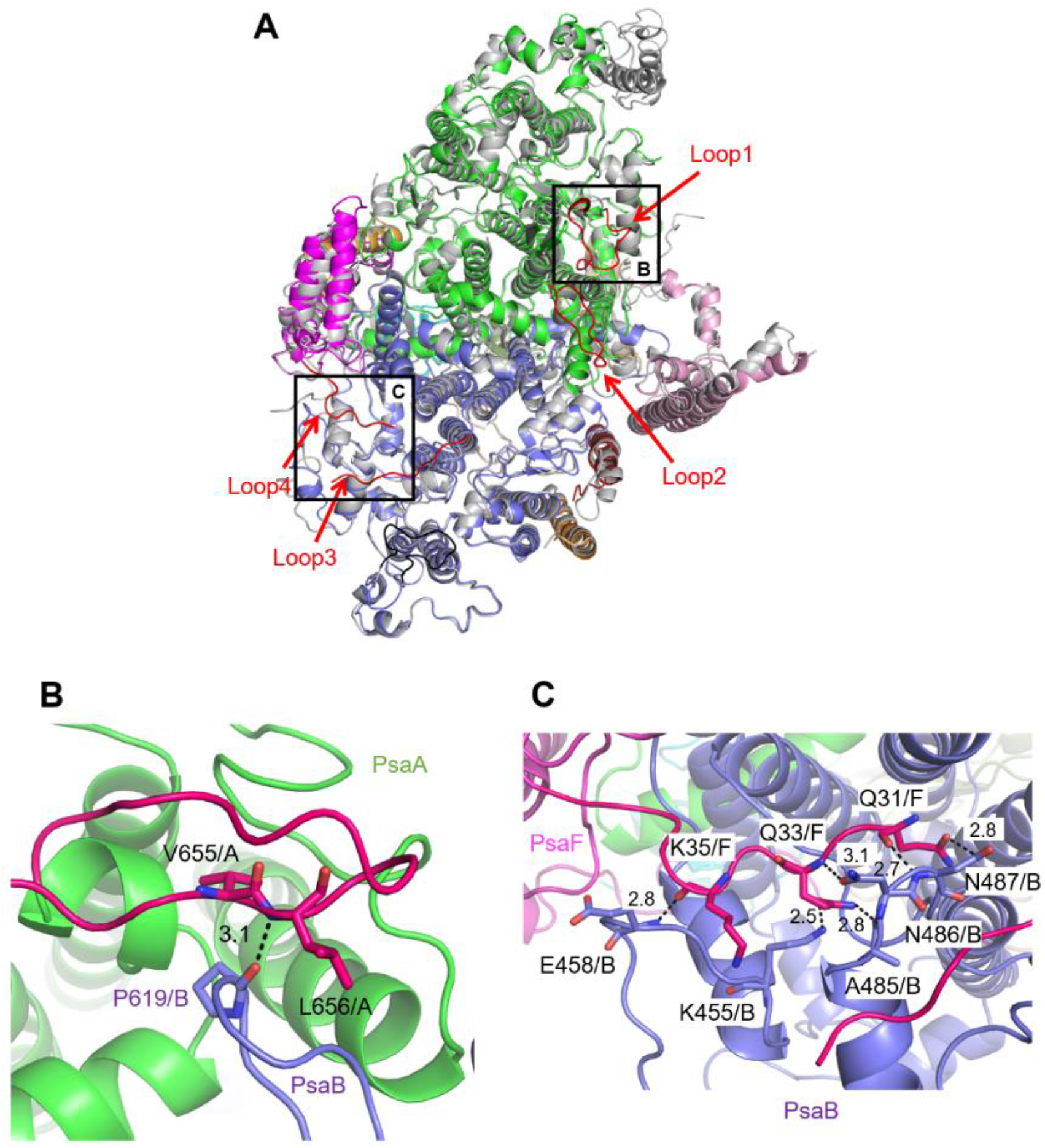
Characteristic structure of the *Gloeobacter* PSI. (**A**) Superposition of the *Gloeobacter* PSI (colored) with the *Synechocystis* PSI (gray) viewed along the membrane normal from the periplasmic side. The structural differences of loop insertions are colored in red. The RMSD is 0.82 Å for 2,019 Cα atoms from subunits observed commonly between *Gloeobacter* and *Synechocystis* PSI. (**B**, **C**) Close-up views of the loop insertions in the regions of Asn652–Ser665 (Loop2) of PsaA in the *Gloeobacter* PSI (**B**) and of Gln31–Asp36 (Loop4) of PsaF in the *Gloeobacter* PSI (**C**).

Both Loop1 of PsaA and Loop3 of PsaB do not interact with other subunits, whereas Loop2 of PsaA and Loop4 of PsaF are elongated so that they are able to interact with PsaB. In Loop2, Leu656 of PsaA is hydrogen-bonded with Pro619 of PsaB at a distance of 3.1 Å, and Val655 of PsaA is in hydrophobic interactions with PsaB-Pro619 (Fig. 3B). In Loop4, Gln31, Gln33, and Lys35 of PsaF interact with Lys455, Glu458, Ala485, Asn486, and Asn487 of PsaB at distances of 2.5–3.1 Å (Fig. 3C). Since our *Gloeobacter* structure is solved at a high resolution, these structural differences cannot be ascribed to uncertainties due to lower resolutions, and the structural features of *Gloeobacter* PSI appear to contribute to the stability and assembly of the PSI complex in the membranes lacking thylakoids.

### Cofactors of the PSI monomer

The cofactors identified in the monomer unit of *Gloeobacter* PSI are summarized in Table S2. There are 89 Chls *a*, 20 *β*-carotenes, 3 [4Fe-4S] clusters, 2 menaquinone-4s, and 4 lipid molecules in each monomer. The location of these molecules is similar to that in other cyanobacterial PSI structures (14–19); however, six Chls are less in the *Gloeobacter* PSI than those in the *T. vulcanus* PSI (16) and *Synechocystis* PSI (15). These Chls include one Chl in PsaA (Chl1A), one Chl in PsaF (Chl1F), two Chls in PsaJ (Chl1J, 2J), and two Chls in PsaK (Chl1K, 2K) (Figs. 4*A* and 4*B*). Both Chl1K and Chl2K do not exist in the *Gloeobacter* PSI because of the absence of PsaK in the genome and the structure. Both Chl1J and Chl2J are located in the periphery of PsaJ, whose amino-acid residues cannot be assigned in the *Gloeobacter* PSI structure. The map quality around Chl1F in the *Gloeobacter* PSI is very high (Fig. 4*C*), indicating that Chl1F is potentially lost in the *Gloeobacter* PSI. Chl1A is difficult to bind to the *Gloeobacter* PSI, because the conserved His residue near Chl1A is changed to Phe243 in the *Gloeobacter* PSI (Figs. S4, S7*A*, S7*B*). This causes a steric hindrance between Phe243 and Chl1A in the *Gloeobacter* PSI (Fig. 4*D* and Fig. S7*A*), making the Chl absent. In addition, it is interesting to note that both of the *T. vulcanus* and *H. hongdechloris* PSI possess an extra Chl (Chl1B) in PsaB (16, 17), which is absent in both *Gloeobacter* PSI and *Synechocystis* PSI (15). This is due to the changes occurred in the loop structures (Fig. 4*E* and Fig. S5), as suggested by Toporik et al. (42).

**Fig. 4.**
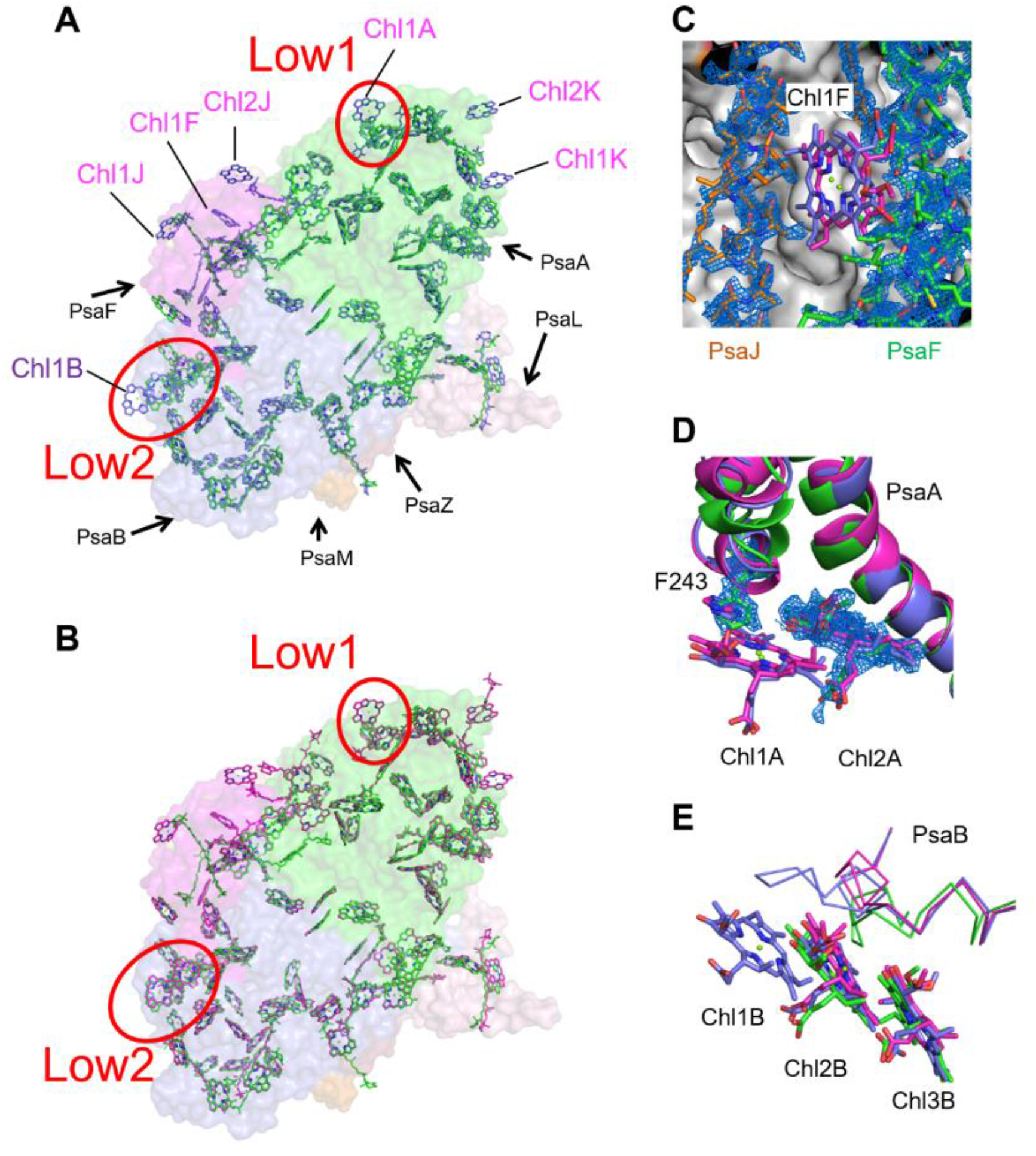
Comparison of pigments among the three types of PSI. (**A**, **B**) Superposition of the PSI monomer viewed along the membrane normal from the periplasmic side, with protein subunits depicted in a surface model. (**A**) *Gloeobacter* (green) vs. *T. vulcanus* (purple); (**B**) *Gloeobacter* (green) vs. *Synechocystis* (magenta). Red circles stand for the sites of Low1 and Low2. (**C**–**E**) Close-up views of nearby pigments of Chl1F (**C**), Chl1A with its densities depicted at 1.5 σ (**D**), and Chl1B (**E**).

### Identification of the sites of Low1 and Low2 in cyanobacterial PSI

Upon comparison of the *Gloeobacter* PSI structure with those of the *Synechocystis* PSI and *T. vulcanus* PSI, we identified two types of low-energy Chls, Low1 and Low2, responsible for the long-wavelength fluorescence peaks as observed in steady-state fluorescence-emission spectra (30–32). The spectrum of *Synechocystis* PSI (red line in Fig. 5*A*) exhibits a fluorescence band at around 723 nm corresponding to Low1, whereas that of *T. vulcanus* PSI (blue line in Fig. 5*A*) displays a fluorescence band at around 731 nm corresponding to Low2. In addition, the width of the fluorescence band from the *T. vulcanus* PSI is broader than that from the *Synechocystis* PSI; therefore, it seems highly probable that the *T. vulcanus* PSI also possesses Low1 in addition to Low2. In contrast, the fluorescence spectrum of *Gloeobacter* PSI (black line in Fig. 5*A*) showed a maximum band at around 694 nm, which is much shorter than the wavelengths of Low1 and Low2. No fluorescence corresponding to Low1 and Low2 are observed in the *Gloeobacter* PSI spectrum, indicating their absence. The characteristic differences of fluorescence peaks in these cyanobacteria can even be observed in absorption spectra measured at 77 K, which showed different intensities of relative absorbance over 700 nm (Fig. 5*B*).

**Fig. 5.**
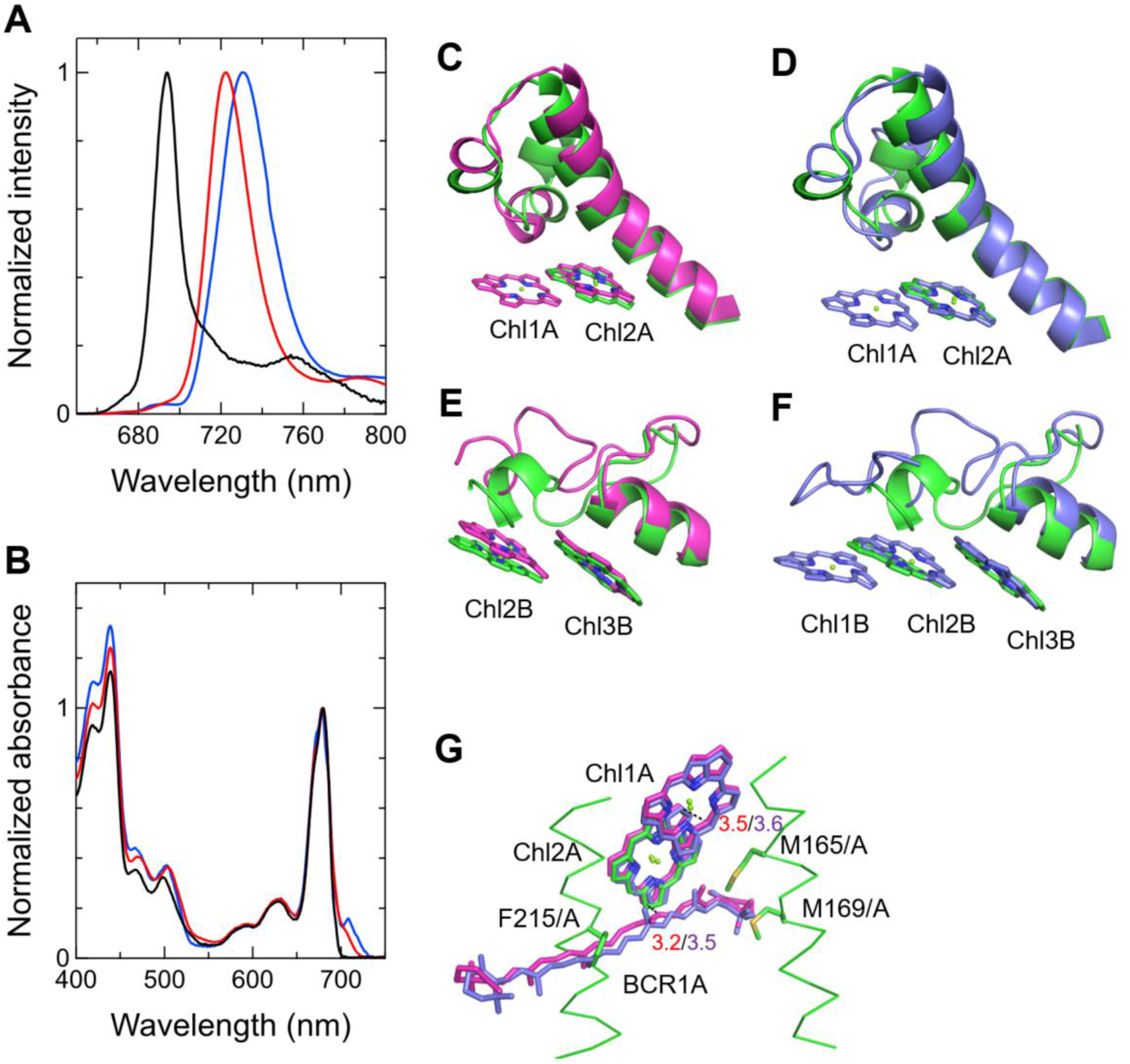
Identification of Low1 and Low2 by spectroscopic properties and structural comparisons. (**A**) Fluorescence spectra of the *Gloeobacter* PSI (black), *Synechocystis* PSI (red), and *T. vulcanus* PSI (blue) measured at 77 K. (**B**) Absorbance spectra of the *Gloeobacter* PSI (black), *Synechocystis* PSI (red), and *T. vulcanus* PSI (blue) measured at 77 K. (**C**) Superposition of the Low1 site in the *Gloeobacter* PSI (green) and *Synechocystis* PSI (magenta). (**D**) Superposition of the Low1 site in the *Gloeobacter* PSI (green) and *T. vulcanus* PSI (purple). (**E**) Superposition of the Low2 site in the *Gloeobacter* PSI (green) and *Synechocystis* PSI (magenta). (**F**) Superposition of the Low2 site in the *Gloeobacter* PSI (green) and *T. vulcanus* PSI (purple). (**G**) Comparison of Low1 with a neighboring *β*-carotene among the three types of cyanobacteria. Green, *Gloeobacter*; magenta, *Synechocystis*; and purple, *T. vulcanus*. Interactions are indicated by dashed lines with distances labeled in Å.

The energy levels of Chls are lowered by formation of dimeric or trimeric Chls; therefore, we focused on dimeric or trimeric Chl clusters through the structural comparisons. Among the six Chl molecules that are absent in the *Gloeobacter* PSI mentioned above, Chl1A forms a dimer with Chl2A in both *Synechocystis* PSI (Fig. 5*C*) and *T. vulcanus* PSI (Fig. 5*D*). These findings indicate that the dimeric Chls of Chl1A/Chl2A is responsible for Low1 in the *Synechocystis* and *T. vulcanus* PSI, which is lost in the *Gloeobacter* PSI due to the loss of Chl1A.

The Mg atom of Chl1A is coordinated by a water molecule in the PSI structures of *T. elongatus* (14) and *H. hongdechloris* (17). In *T. elongatus*, this water molecule is hydrogen-bonded to a neighboring His240 (Fig. S7*A*), indicating that this hydrogen-bond contributes to the stability of Chl1A. Sequence alignment showed that this His240 is conserved among most of cyanobacteria (Fig. S4); however, it is changed to Phe243 in the *Gloeobacter* PsaA (Figs. S4 and S7*A*). As mentioned above, this change leads to the loss of the water molecule that is able to coordinate the Chl molecule (Fig. S7*A*). Thus, both of the His residue and water molecule are responsible for the binding of Chl1A and the formation of Low1.

Low2 exists in the *T. vulcanus* PSI but is absent in the *Gloeobacter* and *Synechocystis* PSI (Figs. 4*A* and 5*A*). Structural comparisons of *T. vulcanus* PSI with *Gloeobacter* PSI and *Synechocystis* PSI showed that a trimeric Chl cluster of Chl1B/Chl2B/Chl3B is present in the *T. vulcanus* PSI but is absent in *Gloeobacter* and *Synechocystis* PSI due to the absence of Chl1B in the latter two cyanobacteria (Figs. 5*E* and 5*F*). This Chl trimer is therefore a most probable candidate for Low2. This is in consistent with the lower energy that Low2 has than Low1, and also supported by the cryo-EM structure of PSI mutant complexes (43).

### Conservation of Low1 and Low2 in PSI-monomer units but not in their interfaces

Some cyanobacteria have tetrameric PSI complexes whose structures have been solved from the cyanobacterium *Anabaena* sp. PCC 7120 (20, 21). Both Low1 and Low2 exist in the monomer units of the *Anabaena* PSI tetramer (Figs. S7*B* and S7*C*). This is in good agreement with the result of the fluorescence spectrum of the *Anabaena* PSI tetramer at 77 K showing a peak at around 730 nm (20, 33). Thus, Low1 and/or Low2 are provided by PSI monomer units from most of the cyanobacteria other than *Gloeobacter*, irrespective of their oligomeric states.

The positions of Low1 and Low2 are found to be within each monomer unit of both PSI trimer and tetramer, but not in the interfaces among the PSI monomers. Historically, low-energy Chls in PSI are thought to be located in interfaces among monomers in a PSI trimer, as summarized briefly by Çoruh et al. (43). This relies on biophysical analyses of various types of PSI monomers isolated from cyanobacteria having PSI trimers. Such interface-located types of low-energy Chls seem to also be observed in *Anabaena* PSI tetramer, which showed a slight shift of fluorescence peak in the PSI dimer and monomer (33). These observations provide evidence for the existence of some low-energy Chls in the interfaces among PSI-monomer units within the PSI oligomers. However, the interface-type low-energy Chls are different from the Low1 and Low2 observed here. Of our findings on the positions of Low1 and Low2, the location of Low2 is supported by the results of Çoruh et al. showing that the PSI monomer, which lacks a Chl molecule B1233 corresponding to Chl1B, has a clear fluorescence peak shift to around 720 nm compared with a fluorescence peak at around 730 nm in the PSI trimer (43).

### Functional significance of Low1 in cyanobacterial PSI

It is known that *Gloeobacter* is susceptible to high-irradiance stress (37, 38), and hence, cannot grow at 30 μmol photons m^−2^ s^−1^. Since Low1 is absent only in the *Gloeobacter* PSI compared with the *Synechocystis* and *T. vulcanus* PSI (Figs. 5*A*, 5*C*, and 5*D*), it seems likely that Low1 is responsible for the light tolerance in *Synechocystis* and *T. vulcanus*. The light tolerance may be induced by excitation-energy quenching by interactions of Chls with Cars (4, 8–10). At the site of Low1, a *β*-carotene molecule BCR1A is located at a distance of 3.2 or 3.5 Å to Chl2A in the *Synechocystis* or *T. vulcanus* PSI structures (Fig. 5*G*). However, the *Gloeobacter* PSI does not possess this *β*-carotene molecule. This is probably due to steric hindrances by side chains of Met165/Met169/Phe215 in the *Gloeobacter* PSI (Fig. 5*G*). These structural differences imply that the tight association of Low1 with the neighboring *β*-carotene serves as a major contributor for the photoprotection machinery of PSI in *Synechocystis* and *T. vulcanus*, which is absent in the primitive cyanobacterium *Gloeobacter*.

Regarding the excitation-energy dynamics in PSI, it is of note that the fluorescence decay-associated spectrum of *Gloeobacter* PSI at 77 K showed no rise component at around 720 nm with a time constant of 32 ps, whereas that of *Synechocystis* PSI exhibited a large rise component at around 720 nm with the same time constant (31). This suggests that Low1 accepts excitation energy at tens of ps in the *Synechocystis* PSI and then some of the energy trapped by Low1 may be transferred to BCR1A. This quenching process may be caused by P700 oxidation as has been suggested previously (25–27).

## Discussion

The overall structures of PSI supercomplexes where the PSI core is associated with their light-harvesting complexes (LHCs) have been solved in red algae (44, 45), green algae (46–49), diatoms (50, 51), moss (52), and higher plants (53, 54). Compared with the structure of *Gloeobacter* PSI (Fig. 6*A*), Low1 is present in the PSI cores from all organisms whose structures have been solved, including a red alga *Cyanidioschyzon merolae* (PDB: 5ZGB), a green alga *Chlamydomonas reinhardtii* (PDB: 6JO5), a diatom *Chaetoceros gracilis* (PDB: 6L4U), a moss *Physcomitrella patens* (PDB: 6L35), and a higher plant *Pisum sativum* (PDB: 4XK8). The His residue corresponding to PsaA-His240 in *T. elongatus* is also conserved among the eukaryotes (Fig. 6*B*), reflecting the presence of Low1 in eukaryotes. Since these organisms grow sufficiently under normal-light conditions such as 30 μmol photons m^−2^ s^−1^, Low1 seems to be an important component of either energy transfer or quenching for cell growth under even relatively low-light conditions in both cyanobacteria and photosynthetic eukaryotes. Thus, Low1 may be responsible for the photoprotection mechanism essential for light-harvesting and survival strategies in oxyphototrophs even under normal-light conditions.

**Fig. 6.**
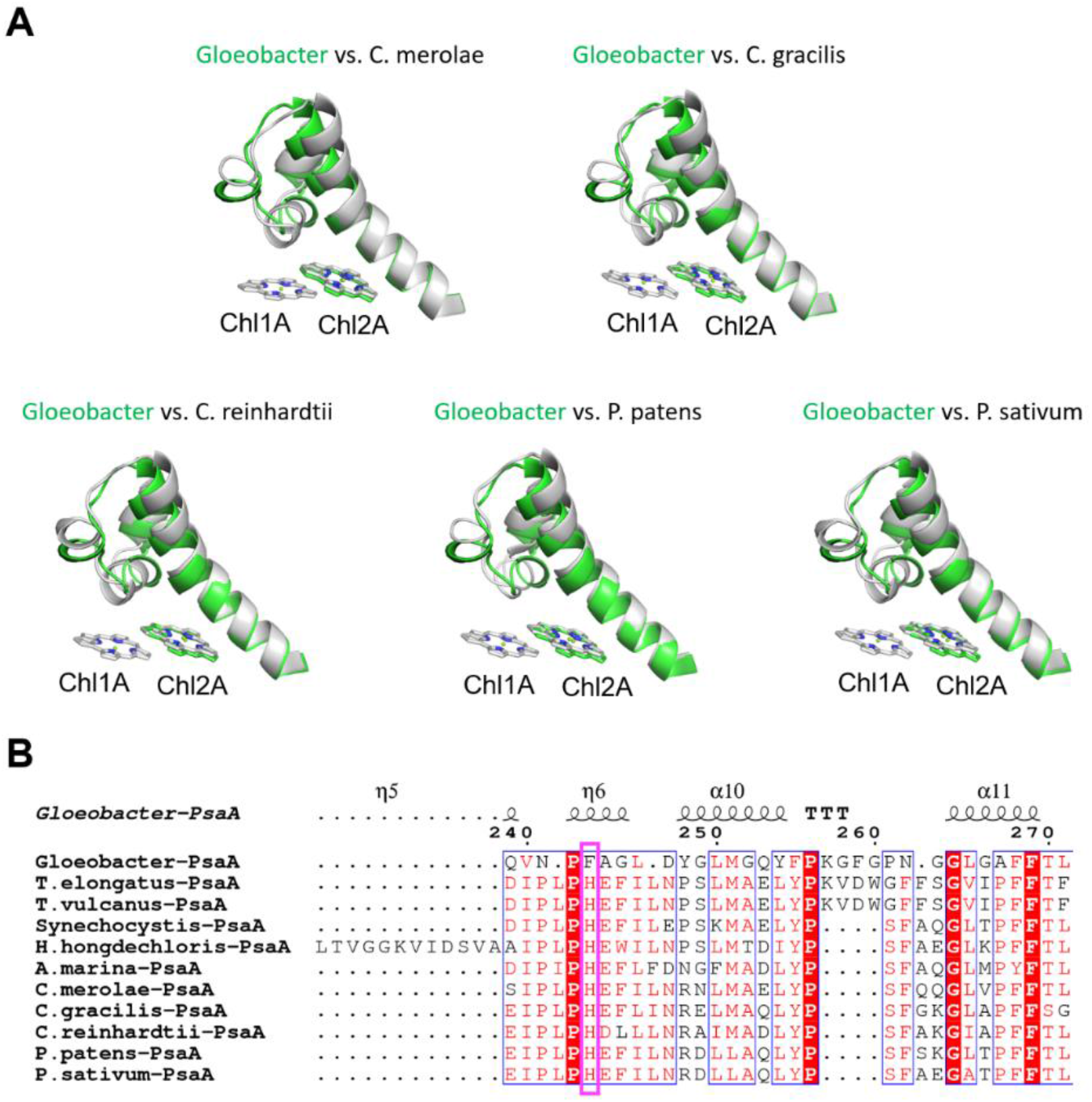
Structural comparisons of the Low1 site among oxyphototrophs. (**A**) Superposition of the Low1 site in the *Gloeobacter* PSI (green) with PSI complexes of photosynthetic eukaryotes (gray) such as *Cyanidioschyzon merolae* (PDB: 5ZGB), *Chaetoceros gracilis* (PDB: 6L4U), *Chlamydomonas reinhardtii* (PDB: 6JO5), *Physcomitrella patens* (PDB: 6L35), and *Pisum sativum* (PDB: 4XK8). (**B**) Multiple sequence alignment of PsaA among oxyphototrophs using ClustalW and ESPript. The species shown are *Gloeobacter violaceus* PCC 7421, *Thermosynechococcus elongatus* BP-1, *Thermosynechococcus vulcanus* NIES-2134, *Synechocystis* sp. PCC 6803, *Halomicronema hongdechloris* C2206, *Acaryochloris marina, Cyanidioschyzon merolae*, *Chaetoceros gracilis*, *Chlamydomonas reinhardtii*, *Physcomitrella patens*, and *Pisum sativum*. The pink box displays the Histidine residue involved in the binding of Chl1A.

Low2 exists only in PSIs of *P. patens* and *P. sativum* in addition to *T. vulcanus* (Fig. 7*A*); however, the triply stacked interactions among three Chls, especially the orientation of Chl1B and Chl1G, differ between cyanobacteria and the eukaryotes (Fig. 7*B*). This is likely due to the different loop structures interacting with Chl1B between *T. vulcanus* and the eukaryotes (Figs. 7*B* and 7*C*). In the *P. sativum* PSI, Chl1G corresponding to Chl1B is bound to Tyr93 of PsaG via a water molecule (Fig. S8*A*). PsaG is present in *C. reinhardtii*; however, the characteristic Tyr residue is substituted to Gly (Fig. S8*B*), leading to the loss of this water molecule in *C. reinhardtii*. This suggests that the hydrogen-bond interaction between the water molecule and Tyr93 of PsaG is required for formation of Low2 in eukaryotes. In contrast, the gene of *psaG* is lost in the red alga and diatom, resulting in the loss of Low2 in these organisms. The different types of Low2 in mosses and higher plants might be necessary for their move from ocean to the land in the process of evolution, in order to adapt to a stronger sunlight on the land. However, it should be noted that Low2 is not needed for cell growth under normal light conditions, because of well growth of *Synechocystis* without Low2. Thus, it is concluded that Low2 is not involved in the universal photoprotection mechanism.

**Fig. 7.**
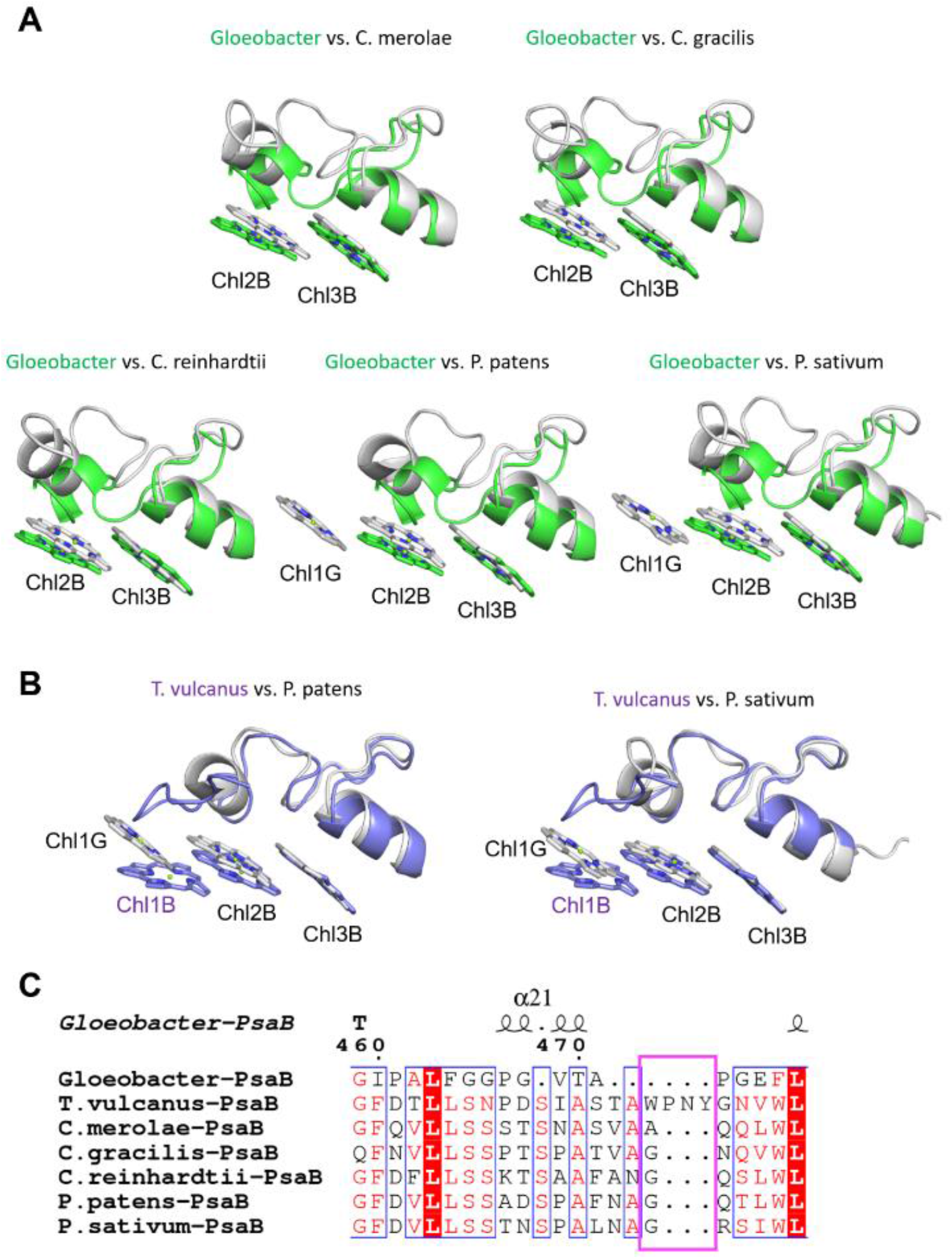
Structural comparisons of the Low2 site among oxyphototrophs. (**A**) Superposition of the Low2 site in the *Gloeobacter* PSI (green) with PSI complexes of photosynthetic eukaryotes (gray) such as *Cyanidioschyzon merolae* (PDB: 5ZGB), *Chaetoceros gracilis* (PDB: 6L4U), *Chlamydomonas reinhardtii* (PDB: 6JO5), *Physcomitrella patens* (PDB: 6L35), and *Pisum sativum* (PDB: 4XK8). (**B**) Superposition of the Low2 site in *T. vulcanus* PSI (purple) with PSI complexes of *P. patens* and *P. sativum* PSI (gray). (**C**) Multiple sequence alignment of PsaB using ClustalW and ESPript. The pink box indicates the loop involved in the binding of Chl1B in *T. vulcanus*.

In conclusion, this study has demonstrated the high-resolution structure of the PSI trimer from *Gloeobacter* at 2.04 Å by cryo-EM. Structural comparisons of *Gloeobacter* PSI with *Synechocystis* and *T. vulcanus* PSI provide evidence for the location of Low1 and Low2 that are responsible for energy quenching in most cyanobacteria but lack in the primitive cyanobacterium *Gloeobacter*. From the physiological behaviors of light susceptibility among the three types of cyanobacteria, Low1 may contribute to a mechanism of photoprotection in PSI by excitation-energy dissipation through Chl-Car interactions, whereas Low2 may be necessary for some organisms, especially land plants, to protect them from higher light intensity. These findings may answer the long-standing question as to the location and function of low-energy Chls in cyanobacterial PSI. Interestingly, the Low1-related photoprotection mechanism may have been conserved in most of the oxyphototrophs other than *Gloeobacter*, suggesting that the lack of Low1 is a characteristic of primitive cyanobacteria occurring near the origin of oxyphototrophs. Thus, this study provides novel insights into not only the mechanism of photoprotection provided by low-energy Chls in PSI, but also the evolution of photosynthetic organisms.

## Materials and Methods

### Purification and characterization of the PSI trimer from *Gloeobacter*

The cyanobacterium *Gloeobacter violaceus* PCC 7421 was grown in BG11 medium (55) supplemented with 10 mM Hepes-KOH (pH 8.0) and 1/1000 volume of KW21 (Daiichi Seimo) at a photosynthetic photon flux density (PPFD) of 5 μmol photons m^−2^ s^−1^ at 20°C with bubbling of air containing 3% (v/v) CO_2_. KW21 is helpful for enhancing the growth of photosynthetic organisms (56, 57). Membrane fragments were prepared after disruption of the cells with glass beads with a method similar to the preparation of thylakoid membranes as described previously (58), and suspended in a buffer containing 0.2 M trehalose, 20 mM Mes-NaOH (pH 6.5), 5 mM CaCl_2_, and 10 mM MgCl_2_ (buffer A). The membranes were solubilized with 1% (w/v) *n*-dodecyl-*β*-D-maltoside (*β*-DDM) at a Chl concentration of 0.25 mg mL^−1^ for 30 min on ice in the dark with gentle stirring. After centrifugation at 50,000 × g for 30 min at 4°C, the resultant supernatant was loaded onto a Q-Sepharose anion-exchange column (2.5 cm of inner diameter and 6 cm of length) equilibrated with buffer A containing 0.03% *β*-DDM (buffer B). The column was washed with buffer B containing 100 mM NaCl (buffer C) until the eluate became colourless, and further washed with 60 mL of buffer B containing 150 mM NaCl. The PSI fraction was eluted with buffer B containing 200 mM NaCl, and subsequently loaded onto a linear trehalose gradient of 10–40% (w/v) in a medium containing 20 mM Mes-NaOH (pH 6.5), 5 mM CaCl_2_, 10 mM MgCl_2_, 100 mM NaCl, and 0.03% *β*-DDM. After centrifugation at 154,000 × g for 18 h at 4°C (P40ST rotor; Hitachi), a band containing the PSI trimers was collected and then concentrated using a 150 kDa cut-off filter (Apollo; Orbital Biosciences) at 4,000 × g. The concentrated PSI core trimer complexes were stored in liquid nitrogen until use.

Subunit composition of the PSI was analyzed by SDS-polyacrylamide gel electrophoresis (PAGE) containing 16% acrylamide and 7.5 M urea according to a method by Ikeuchi and Inoue (59) (Fig. 1*A*). The samples (5 μg of Chl) were solubilized by 3% lithium lauryl sulfate and 75 mM dithiothreitol for 10 min at 60°C, and loaded onto the gel. A standard molecular weight marker (SP-0110; APRO Science) was used. The subunit bands were assigned by mass spectrometry according to a previous method (60). Pigment compositions were analysed as described in Nagao et al. (61), and the elution profile was monitored at 440 nm (Fig. S1*B*).

### Purification of the PSI trimer from *Synechocystis*

The cyanobacterium *Synechocystis* sp. PCC 6803 47-H strain was grown in the BG11 medium supplemented with both 10 mM Hepes-KOH (pH 8.0) at the PPFD of 30 μmol photons m^−2^ s^−1^ at 30°C with bubbling of air containing 3% (v/v) CO_2_. The 47-H strain has a six-histidine tag at the C-terminus of the CP47 subunit (62). Thylakoid membranes were prepared (58) and suspended in buffer A. The thylakoids were solubilized with 1% *β*-DDM at a Chl concentration of 0.50 mg mL^−1^ for 30 min on ice in the dark with gentle stirring. After centrifugation at 50,000 × g for 10 min at 4°C, the resultant supernatant was loaded onto a Ni^2+^ affinity column (2.5 cm of inner diameter and 10 cm of length) equilibrated with buffer C. The PSI-enriched fraction was collected by washing the column with buffer C, and subsequently diluted with an equal volume of buffer B. The diluted sample was applied onto a Q-Sepharose anion-exchange column (2.5 cm of inner diameter and 10 cm of length) equilibrated with buffer B. The column was washed with buffer B containing 150 mM NaCl until the eluate became colourless, and further washed with 50 mL of buffer B containing 200 mM NaCl and subsequently with 100 mL of buffer B containing 250 mM NaCl. The PSI fraction was eluted with buffer B containing 300 mM NaCl, and subsequently loaded onto a linear trehalose gradient of 10–40% (w/v) in a medium containing 20 mM Mes-NaOH (pH 6.5), 5 mM CaCl_2_, 10 mM MgCl_2_, 100 mM NaCl, and 0.03% *β*-DDM. After centrifugation at 154,000 × g for 18 h at 4°C (P40ST rotor; Hitachi), a band containing the PSI trimers was collected and then concentrated using a 150 kDa cut-off filter (Apollo; Orbital Biosciences) at 4,000 × g. The concentrated PSI core trimer complexes were stored in liquid nitrogen until use.

### Purification of the PSI trimer from *T. vulcanus*

The cyanobacterium *Thermosynechococcus vulcanus* NIES-2134 was grown and its thylakoid membranes were prepared as described previously (63). The PSI-enriched fraction was obtained as precipitation by centrifugation at 100,000 × g for 60 min at 4°C after second-round treatments of the thylakoids with *N*, *N*-dimethyldodecylamine-*N*-oxide (63), and suspended with a 30 mM Mes-NaOH (pH 6.0) buffer containing 5% (w/v) glycerol, 3 mM CaCl_2_, and 0.03% *β*-DDM. The PSI fraction was loaded onto a linear trehalose gradient of 10–40% (w/v) in a medium containing 20 mM Mes-NaOH (pH 6.5), 5 mM CaCl_2_, 10 mM MgCl_2_, 100 mM NaCl, and 0.03% *β*-DDM. After centrifugation at 154,000 × g for 18 h at 4°C (P40ST rotor; Hitachi), a band containing the PSI trimers was collected and then concentrated using a 150 kDa cut-off filter (Apollo; Orbital Biosciences) at 4,000 × g. The concentrated PSI core trimer complexes were stored in liquid nitrogen until use.

### Absorption and fluorescence-emission spectra at 77 K

Absorption spectra of the three types of PSI cores were measured at 77 K using a spectrometer equipped with an integrating sphere unit (V-650/ISVC-747, JASCO) (64), and their steady-state fluorescence-emission spectra were recorded at 77 K using a spectrofluorometer (FP-8300/PMU-183, JASCO). Excitation wavelength was set to 440 nm.

### Cryo-EM data collection

For cryo-EM experiments, 3-μL aliquots of the *Gloeobacter* PSI trimers (1.68 mg Chl mL^−1^) were applied to Quantifoil R1.2/1.3, Cu 200 mesh grids pre-treated by gold sputtering. Without waiting for incubation, the excess amount of the solution was blotted off for 5 sec with a filter paper in an FEI Vitrobot Mark IV at 4°C under 100% humidity. The grids were plunged into liquid ethane cooled by liquid nitrogen and then transferred into a CRYO ARM 300 electron microscope (JEOL) equipped with a cold-field emission gun operated at 300 kV. Zero-energy loss images were recorded at a nominal magnification of × 60,000 on a direct electron detection camera (Gatan K3, AMETEK) with a nominal defocus range of −1.8 to −0.6 μm. One-pixel size corresponded to 0.823 Å. Each image stack was exposed at a dose rate of 17.555 e^−^Å^−2^sec^−1^ for 4.0 sec, and consisted of dose-fractionated 50 movie frames. In total 7,282 image stacks were collected.

### Cryo-EM image processing

The movie frames thus obtained were aligned and summed using MotionCor2 (65) to yield dose weighted images. Estimation of the contrast transfer function (CTF) was performed using CTFFIND4 (66). All of the following processes were performed using RELION3.1 (67). In total 961,960 particles were automatically picked up and used for reference-free 2D classification. Then, 946,827 particles were selected from good 2D classes and subsequently subjected to 3D classification without imposing any symmetry. An initial model for the first 3D classification was generated *de novo* from 2D classifications. As shown in Fig. S2, the final PSI trimer structure was reconstructed from 261,743 particles. The overall resolution of the cryo-EM map was estimated to be 2.04 Å by the gold-standard FSC curve with a cut-off value of 0.143 (Fig. S2*D*) (68). Local resolutions were calculated using RELION (Fig. S2*F*).

### Model building and refinement

The cryo-EM map thus obtained was used for model building of the PSI trimer. Each subunit of the homology models constructed using the Phyre2 server (69) was first manually fitted into the map by using UCSF Chimera (70), and then inspected and manually adjusted with Coot (71). The complete PSI-trimer structure was refined with phenix.real_space_refine (72) and REFMAC5 (73) with geometric restraints for the protein–cofactor coordination. The final model was validated with MolProbity (74), EMringer (75), and Q-score (76). The statistics for all data collection and structure refinement are summarized in Tables S1 and S2. All structural figures are made by Pymol (77) or UCSF ChimeraX (78).

## Data availability

Atomic coordinates and cryo-EM maps for the reported structure of the PSI trimer have been deposited in the Protein Data Bank under an accession code 7F4V and in the Electron Microscopy Data Bank under an accession code EMD-31455.

## Acknowledgments

This work was supported by JSPS KAKENHI grant Nos. JP20H02914 (Koji.K.), JP21K19085 (R.N.), JP20K06528 (KeisukeK), JP16H06553 (S.A.), and JP17H06433 (J.-R.S.), JST-Mirai Program Grant Number JPMJMI20G5 (K.Y.), and the Cyclic Innovation for Clinical Empowerment (CiCLE) from the Japan Agency for Medical Research and Development, AMED (T.H., Keisuke.K., K.Y.).

## Author Contributions

R.N. and J.-R.S. conceived the project; H.U. and A.M. prepared the stabilized culture stain of *Gloeobacter;* R.N. and Y.N. cultured *Gloeobacter, Synechocystis*, and *T. vulcanus;* R.N. purified the PSI-trimer complexes from the three types of cyanobacteria, and characterized their biochemical properties; Y.U., M.Y., and S.A. performed spectroscopic measurements of the three types of PSI trimers and their data analysis; T.S. and N.D. identified the PSI subunits of *Gloeobacter* by mass spectrometry; T.H. collected cryo-EM images; Koji K. processed the EM data and reconstructed the final EM map; Koji K. built the structural model and refined the final model; Keisuke K. commented on the structural data; K.Y. and J.-R.S. supervised this project; Koji K. and R.N. wrote the draft manuscript; and R.N. and J.-R.S. wrote the final manuscript, and all of the authors joined the discussion of the results.

## References

1. R. E. Blankenship, Molecular Mechanisms of Photosynthesis (Wiley-Blackwell, Oxford, U.K., ed. 3rd, 2021).

2. M. Edelman, A. K. Mattoo, D1-protein dynamics in photosystem II: the lingering enigma. Photosynth. Res. 98, 609–620 (2008).

3. N. Murata, S. I. Allakhverdiev, Y. Nishiyama, The mechanism of photoinhibition *in vivo:* Re-evaluation of the roles of catalase, α-tocopherol, non-photochemical quenching, and electron transport. Biochim. Biophys. Acta, Bioenerg. 1817, 1127–1133 (2012).

4. A. V. Ruban, M. P. Johnson, C. D. P. Duffy, The photoprotective molecular switch in the photosystem II antenna. Biochim. Biophys. Acta, Bioenerg. 1817, 167–181 (2012).

5. P. Müller, X.-P. Li, K. K. Niyogi, Non-photochemical quenching. A response to excess light energy. Plant Physiol. 125, 1558–1566 (2001).

6. P. Horton, A. Ruban, Molecular design of the photosystem II light-harvesting antenna: Photosynthesis and photoprotection. J. Exp. Bot. 56, 365–373 (2005).

7. P. Jahns, A. R. Holzwarth, The role of the xanthophyll cycle and of lutein in photoprotection of photosystem II. Biochim. Biophys. Acta, Bioenerg. 1817, 182–193 (2012).

8. Y.-Z. Ma, N. E. Holt, X.-P. Li, K. K. Niyogi, G. R. Fleming, Evidence for direct carotenoid involvement in the regulation of photosynthetic light harvesting. Proc. Natl. Acad. Sci. U. S. A. 100, 4377–4382 (2003).

9. A. V. Ruban et al., Identification of a mechanism of photoprotective energy dissipation in higher plants. Nature 450, 575–578 (2007).

10. T. K. Ahn et al., Architecture of a charge-transfer state regulating light harvesting in a plant antenna protein. Science 320, 794–797 (2008).

11. K. Brettel, W. Leibl, Electron transfer in photosystem I. Biochim. Biophys. Acta, Bioenerg. 1507, 100–114 (2001).

12. N. Nelson, C. F. Yocum, Structure and function of photosystems I and II. Annu. Rev. Plant Biol. 57, 521–565 (2006).

13. M. Suga, J.-R. Shen, Structural variations of photosystem I-antenna supercomplex in response to adaptations to different light environments. Curr. Opin. Struct. Biol. 63, 10–17 (2020).

14. P. Jordan et al., Three-dimensional structure of cyanobacterial photosystem I at 2.5 Å resolution. Nature 411, 909–917 (2001).

15. T. Malavath, I. Caspy, S. Y. Netzer-El, D. Klaiman, N. Nelson, Structure and function of wild-type and subunit-depleted photosystem I in *Synechocystis*. Biochim. Biophys. Acta, Bioenerg. 1859, 645–654 (2018).

16. F. Akita et al., Structure of a cyanobacterial photosystem I surrounded by octadecameric IsiA antenna proteins. Commun. Biol. 3, 232 (2020).

17. K. Kato et al., Structural basis for the adaptation and function of chlorophyll f in photosystem I. Nat. Commun. 11, 238 (2020).

18. T. Hamaguchi et al., Structure of the far-red light utilizing photosystem I of *Acaryochloris marina*. Nat. Commun. 12, 2333 (2021).

19. C. Xu et al., A unique photosystem I reaction center from a chlorophyll *d*-containing cyanobacterium *Acaryochloris marina*. J. Integr. Plant Biol. DOI: 10.1111/jipb.13113 (2021).

20. K. Kato et al., Structure of a cyanobacterial photosystem I tetramer revealed by cryo-electron microscopy. Nat. Commun. 10, 4929 (2019).

21. L. Zheng et al., Structural and functional insights into the tetrameric photosystem I from heterocyst-forming cyanobacteria. Nat. Plants 5, 1087–1097 (2019).

22. M. Chen et al., Distinct structural modulation of photosystem I and lipid environment stabilizes its tetrameric assembly. Nat. Plants 6, 314–320 (2020).

23. B. Gobets, R. van Grondelle, Energy transfer and trapping in photosystem I. Biochim. Biophys. Acta, Bioenerg. 1507, 80–99 (2001).

24. N. V. Karapetyan, E. Schlodder, R. van Grondelle, J. P. Dekker “The long wavelength chlorophylls of Photosystem I” in Photosystem I: the Light-Driven Plastocyanin:Ferredoxin Oxidoreductase, J. H. Golbeck, Ed. (Springer, Dordrecht, The Netherlands, 2006), pp. 177–192.

25. V. V. Shubin, I. N. Bezsmertnaya, N. V. Karapetyan, Efficient energy transfer from the long-wavelength antenna chlorophylls to P700 in photosystem I complexes from *Spirulina platensis*. J. Photochem. Photobiol. B 30, 153–160 (1995).

26. Y. Shibata, A. Yamagishi, S. Kawamoto, T. Noji, S. Itoh, Kinetically distinct three red chlorophylls in photosystem I of *Thermosynechococcus elongatus* revealed by femtosecond time-resolved fluorescence spectroscopy at 15 K. J. Phys. Chem. B 114, 2954–2963 (2010).

27. E. Schlodder, M. Hussels, M. Çetin, N. V. Karapetyan, M. Brecht, Fluorescence of the various red antenna states in photosystem I complexes from cyanobacteria is affected differently by the redox state of P700. Biochim. Biophys. Acta, Bioenerg. 1807, 1423–1431 (2011).

28. M. Byrdin et al., Light harvesting in photosystem I: modeling based on the 2.5-A structure of photosystem I from *Synechococcus elongatus*. Biophys J. 83, 433–457 (2002).

29. J. Adolphs, F. Müh, M. E.-A. Madjet, M. S. am Busch, T. Renger, Structure-based calculations of optical spectra of photosystem I suggest an asymmetric light-harvesting process. J. Am. Chem. Soc. 132, 3331–3343 (2010).

30. J. van der Lee et al., Steady-state polarized light spectroscopy of isolated Photosystem I complexes. Photosynth. Res. 35, 311–321 (1993).

31. M. Mimuro, M. Yokono, S. Akimoto, Variations in photosystem I properties in the primordial cyanobacterium *Gloeobacter violaceus* PCC 7421. Photochem. Photobiol. 86, 62–69 (2010).

32. L.-O. Pålsson, J. P. Dekker, E. Schlodder, R. Monshouwer, R. van Grondelle, Polarized site-selective fluorescence spectroscopy of the long-wavelength emitting chlorophylls in isolated Photosystem I particles of *Synechococcus elongatus*. Photosynth. Res. 48, 239–246 (1996).

33. R. Nagao et al., pH-induced regulation of excitation energy transfer in the cyanobacterial photosystem I tetramer. J. Phys. Chem. B 124, 1949–1954 (2020).

34. F. Koenig, M. Schmidt, *Gloeobacter violaceus* - investigation of an unusual photosynthetic apparatus. Absence of the long-wavelength emission of photosystem I in 77 K fluorescence spectra. Physiol. Plant. 94, 621–628 (1995).

35. G. Guglielmi, G. Cohen-Bazire, D. A. Bryant, The structure of *Gloeobacter violaceus* and its phycobilisomes. Arch. Microbiol. 129, 181–189 (1981).

36. B. Nelissen, Y. Van de Peer, A. Wilmotte, R. De Wachter, An early origin of plastids within the cyanobacterial divergence is suggested by evolutionary trees based on complete 16S rRNA sequences. Mol. Biol. Evol. 12, 1166–1173 (1995).

37. M. Mimuro et al. (2001) Photosynthetic properties of a cyanobacterium, *Gloeobacter violaceus* PCC7421. in Proceedings of the 12th International Congress on Photosynthesis Brisbane.

38. C. I. Sicora, C. M. Brown, O. Cheregi, I. Vass, D. A. Campbell, The *psbA* gene family responds differentially to light and UVB stress in *Gloeobacter violaceus* PCC 7421, a deeply divergent cyanobacterium. Biochim. Biophys. Acta, Bioenerg. 1777, 130–139 (2008).

39. T. Hamaguchi et al., A new cryo-EM system for single particle analysis. J. Struct. Biol. 207, 40–48 (2019).

40. Y. Nakamura et al., Complete genome structure of *Gloeobacter violaceus* PCC 7421, a cyanobacterium that lacks thylakoids. DNA Res. 10, 137–145 (2003).

41. H. Inoue et al., Unique constitution of photosystem I with a novel subunit in the cyanobacterium *Gloeobacter violaceus* PCC 7421. FEBS Lett. 578, 275–279 (2004).

42. H. Toporik et al., The structure of a red-shifted photosystem I reveals a red site in the core antenna. Nat. Commun. 11, 5279 (2020).

43. O. Çoruh et al., Cryo-EM structure of a functional monomeric Photosystem I from *Thermosynechococcus elongatus* reveals red chlorophyll cluster. Commun. Biol. 4, 304 (2021).

44. X. Pi et al., Unique organization of photosystem I-light-harvesting supercomplex revealed by cryo-EM from a red alga. Proc. Natl. Acad. Sci. U. S. A. 115, 4423–4428 (2018).

45. M. Antoshvili, I. Caspy, M. Hippler, N. Nelson, Structure and function of photosystem I in *Cyanidioschyzon merolae*. Photosynth. Res. 139, 499–508 (2019).

46. X. Su et al., Antenna arrangement and energy transfer pathways of a green algal photosystem-I-LHCI supercomplex. Nat. Plants 5, 273–281 (2019).

47. M. Suga et al., Structure of the green algal photosystem I supercomplex with a decameric light-harvesting complex I. Nat. Plants 5, 626–636 (2019).

48. X. Qin et al., Structure of a green algal photosystem I in complex with a large number of light-harvesting complex I subunits. Nat. Plants 5, 263–272 (2019).

49. A. Perez-Boerema et al., Structure of a minimal photosystem I from the green alga *Dunaliella salina*. Nat. Plants 6, 321–327 (2020).

50. R. Nagao et al., Structural basis for assembly and function of a diatom photosystem I-light-harvesting supercomplex. Nat. Commun. 11, 2481 (2020).

51. C. Xu et al., Structural basis for energy transfer in a huge diatom PSI-FCPI supercomplex. Nat. Commun. 11, 5081 (2020).

52. Q. Yan et al., Antenna arrangement and energy-transfer pathways of PSI-LHCI from the moss *Physcomitrella patens*. Cell Discov. 7, 10 (2021).

53. X. Qin, M. Suga, T. Kuang, J.-R. Shen, Structural basis for energy transfer pathways in the plant PSI-LHCI supercomplex. Science 348, 989–995 (2015).

54. Y. Mazor, A. Borovikova, I. Caspy, N. Nelson, Structure of the plant photosystem I supercomplex at 2.6 Å resolution. Nat. Plants 3, 17014 (2017).

55. R. Y. Stanier, R. Kunisawa, M. Mandel, G. Cohen-Bazire, Purification and properties of unicellular blue-green algae (order Chroococcales). Bacteriol. Rev. 35, 171–205 (1971).

56. R. Nagao et al., Isolation and characterization of oxygen-evolving thylakoid membranes and Photosystem II particles from a marine diatom *Chaetoceros gracilis*. Biochim. Biophys. Acta, Bioenerg. 1767, 1353–1362 (2007).

57. R. Nagao et al., Comparison of oligomeric states and polypeptide compositions of fucoxanthin chlorophyll *a*/*c*-binding protein complexes among various diatom species. Photosynth. Res. 117, 281–288 (2013).

58. R. Nagao, M. Yamaguchi, S. Nakamura, H. Ueoka-Nakanishi, T. Noguchi, Genetically introduced hydrogen bond interactions reveal an asymmetric charge distribution on the radical cation of the special-pair chlorophyll P680. J. Biol. Chem. 292, 7474–7486 (2017).

59. M. Ikeuchi, Y. Inoue, A new photosystem II reaction center component (4.8 kDa protein) encoded by chloroplast genome. FEBS Lett. 241, 99–104 (1988).

60. R. Nagao et al., Structural basis for energy harvesting and dissipation in a diatom PSII-FCPII supercomplex. Nat. Plants 5, 890–901 (2019).

61. R. Nagao, M. Yokono, S. Akimoto, T. Tomo, High excitation energy quenching in fucoxanthin chlorophyll *a*/*c*-binding protein complexes from the diatom *Chaetoceros gracilis*. J. Phys. Chem. B 117, 6888–6895 (2013).

62. H. Ito, A. Tanaka, Evolution of a divinyl chlorophyll-based photosystem in *Prochlorococcus*. Proc. Natl. Acad. Sci. U. S. A. 108, 18014–18019 (2011).

63. K. Kawakami, J.-R. Shen, Purification of fully active and crystallizable photosystem II from thermophilic cyanobacteria. Methods Enzymol. 613, 1–16 (2018).

64. F. Hamada, M. Yokono, E. Hirose, A. Murakami, S. Akimoto, Excitation energy relaxation in a symbiotic cyanobacterium, *Prochloron didemni*, occurring in coral-reef ascidians, and in a free-living cyanobacterium, *Prochlorothrix hollandica*. Biochim. Biophys. Acta, Bioenerg. 1817, 1992–1997 (2012).

65. S. Q. Zheng et al., MotionCor2: anisotropic correction of beam-induced motion for improved cryo-electron microscopy. Nat. Methods 14, 331–332 (2017).

66. J. A. Mindell, N. Grigorieff, Accurate determination of local defocus and specimen tilt in electron microscopy. J. Struct. Biol. 142, 334–347 (2003).

67. J. Zivanov, T. Nakane, S. H. W. Scheres, Estimation of high-order aberrations and anisotropic magnification from cryo-EM data sets in *RELION-3.1*. IUCrJ 7, 253–267 (2020).

68. N. Grigorieff, S. C. Harrison, Near-atomic resolution reconstructions of icosahedral viruses from electron cryo-microscopy. Curr. Opin. Struc. Biol. 21, 265–273 (2011).

69. L. A. Kelley, S. Mezulis, C. M. Yates, M. N. Wass, M. J. E. Sternberg, The Phyre2 web portal for protein modeling, prediction and analysis. Nat. Protoc. 10, 845–858 (2015).

70. E. F. Pettersen et al., UCSF chimera - A visualization system for exploratory research and analysis. J. Comput. Chem. 25, 1605–1612 (2004).

71. P. Emsley, B. Lohkamp, W. G. Scott, K. Cowtan, Features and development of *Coot*. Acta Crystallogr. D Biol. Crystallogr. 66, 486–501 (2010).

72. P. D. Adams et al., PHENIX: a comprehensive Python-based system for macromolecular structure solution. Acta Crystallogr. D Biol. Crystallogr. 66, 213–221 (2010).

73. G. N. Murshudov et al., REFMAC5 for the refinement of macromolecular crystal structures. Acta Crystallogr. D Biol. Crystallogr. 67, 355–367 (2011).

74. V. B. Chen et al., MolProbity: all-atom structure validation for macromolecular crystallography. Acta Crystallogr. D Biol. Crystallogr. 66, 12–21 (2010).

75. B. A. Barad et al., EMRinger: side chain-directed model and map validation for 3D cryo-electron microscopy. Nat. Methods 12, 943–946 (2015).

76. G. Pintilie et al., Measurement of atom resolvability in cryo-EM maps with *Q*-scores. Nat. Methods 17, 328–334 (2020).

77. L. L. C. Schrödinger, The PyMOL Molecular Graphics System. Version 2.4.1. (2021).

78. T. D. Goddard et al., UCSF ChimeraX: Meeting modern challenges in visualization and analysis. Protein Sci. 27, 14–25 (2018).

