## SupplementaryInformation for "Structural basis for the absence of low-energy chlorophylls responsible for photoprotection from a primitive cyanobacterial PSI"

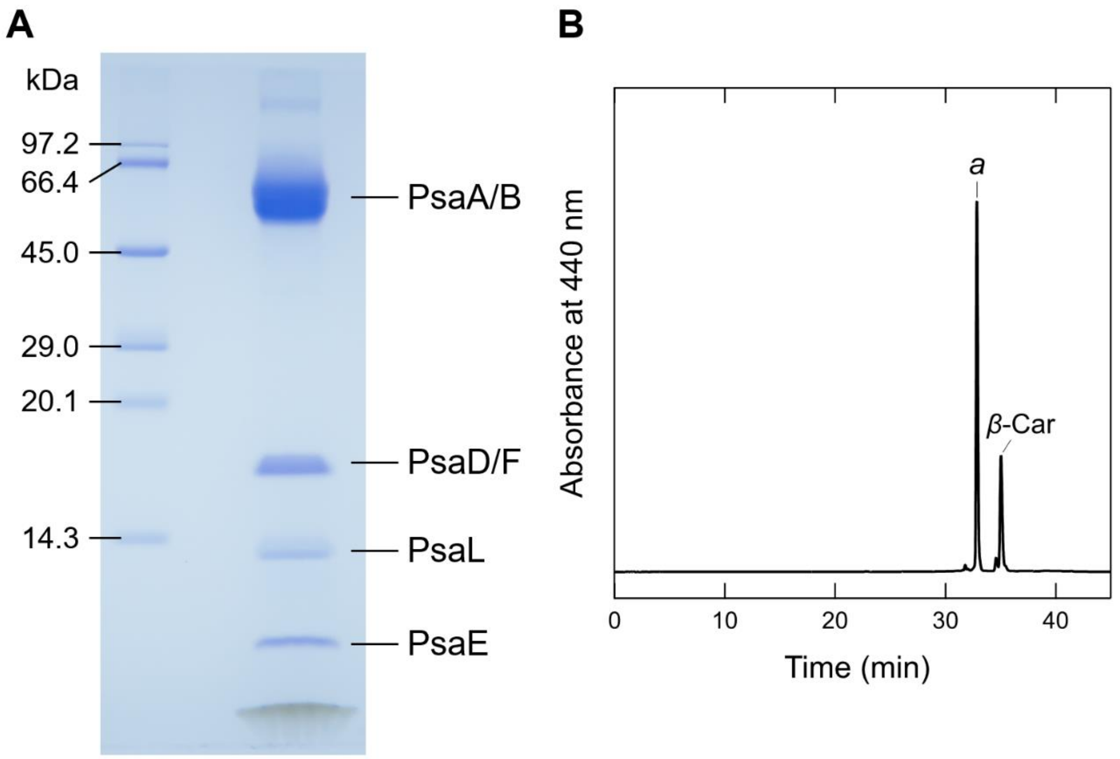


**Fig. S1 Biochemical characterization of the *Gloeobacter* PSI trimer.** (**A**) SDS-PAGE analysis of the PSI trimer. Protein bands were identified by mass spectrometry. It should be noted that some of the subunits in the PSI trimer were not detected by SDS-PAGE probably due to their poor staining by CBB, but were visualized by the present cryo-EM map. (**B**) HPLC analysis of the pigments extracted from PSI monitored at 440 nm. Letters of *a* and *β*-Car indicate Chl *a* and *β*-carotene, respectively.

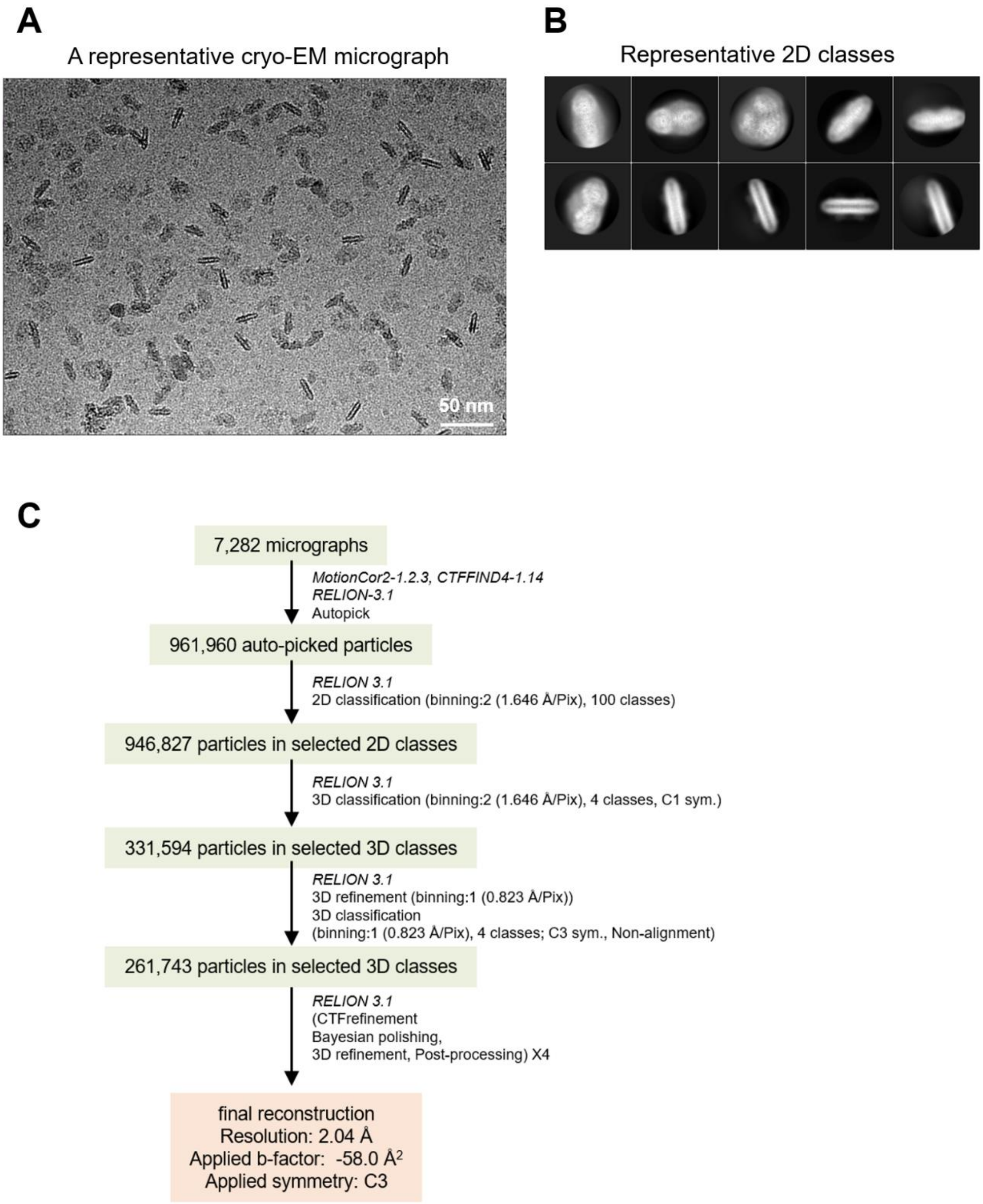

A
A representative cryo-EM micrograph
50 nm
B
Representative 2D classes
C
7,282 micrographs
MotionCor2-1.2.3, CTFFIND4-1.14
RELION-3.1
Autopick
961,960 auto-picked particles
RELION 3.1
2D classification (binning:2 (1.646 Å/Pix), 100 classes)
946,827 particles in selected 2D classes
RELION 3.1
3D classification (binning:2 (1.646 Å/Pix), 4 classes, C1 sym.)
331,594 particles in selected 3D classes
RELION 3.1
3D refinement (binning:1 (0.823 Å/Pix))
3D classification
(binning:1 (0.823 Å/Pix), 4 classes; C3 sym., Non-alignment)
261,743 particles in selected 3D classes
RELION 3.1
(CTFrefinement
Bayesian polishing,
3D refinement, Post-processing) X4
final reconstruction
Resolution: 2.04 Å
Applied b-factor: -58.0 Å2
Applied symmetry: C3

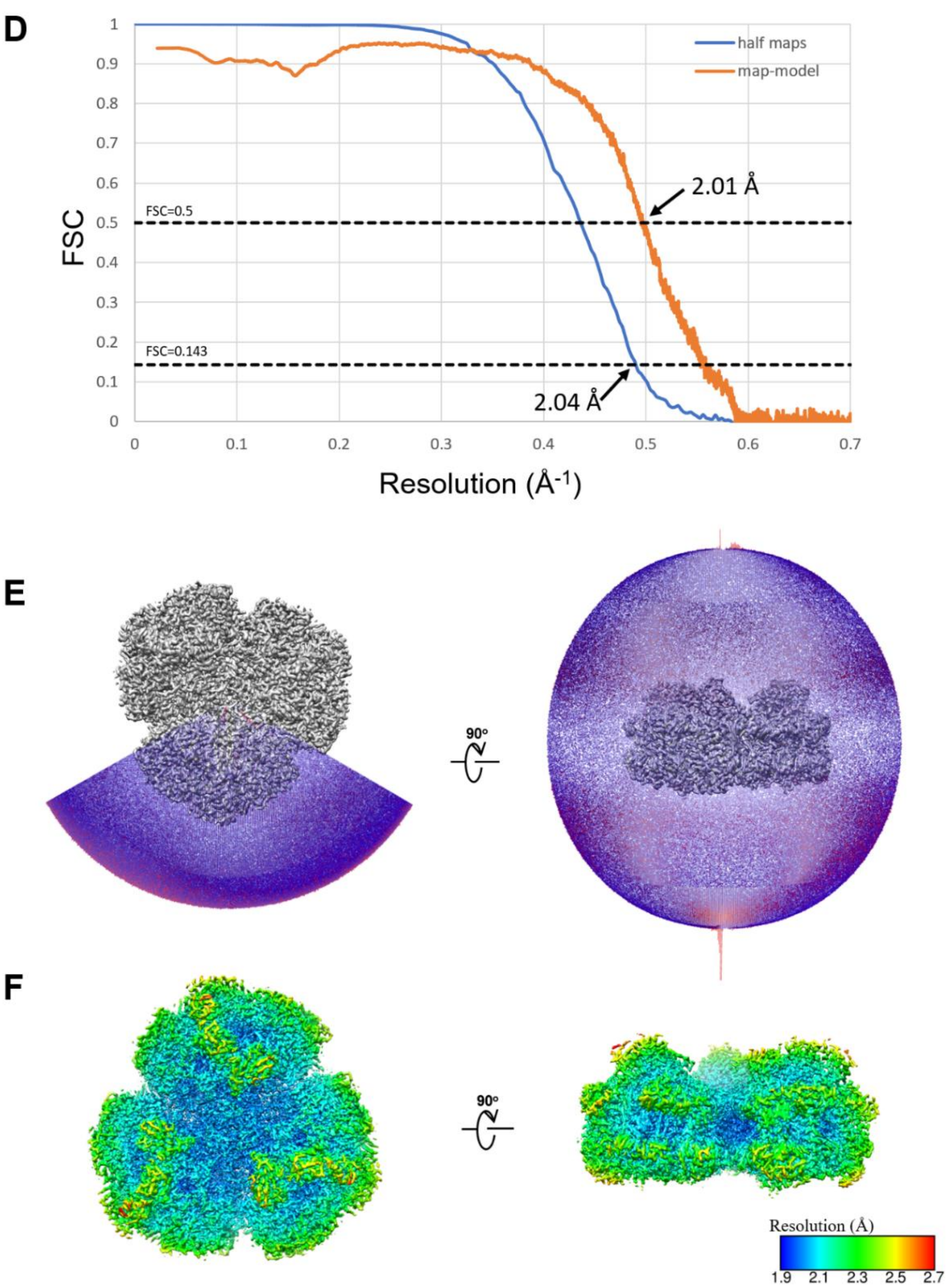


**Fig. S2 Cryo-EM data collection and processing of the *Gloeobacter* PSI trimer.** (**A**) A representative cryo-EM micrograph of the PSI trimer. (**B**) Representative 2D classes of the PSI trimer. The box size is 330 Å. (**C**) A flowchart showing the classification scheme of the PSI trimer. The overall structure of PSI was reconstructed at a 2.04-Å resolution from 261,743 particles. (**D**) FSC curves of the PSI trimer for independently refined half-maps (half maps) and full map vs. model (map-model). (**E**) Angular distribution of the particles used for reconstruction of the PSI trimer. Each cylinder represents one view, and its height is proportional to the number of particles. (**F**) Local resolution map of the PSI trimer.

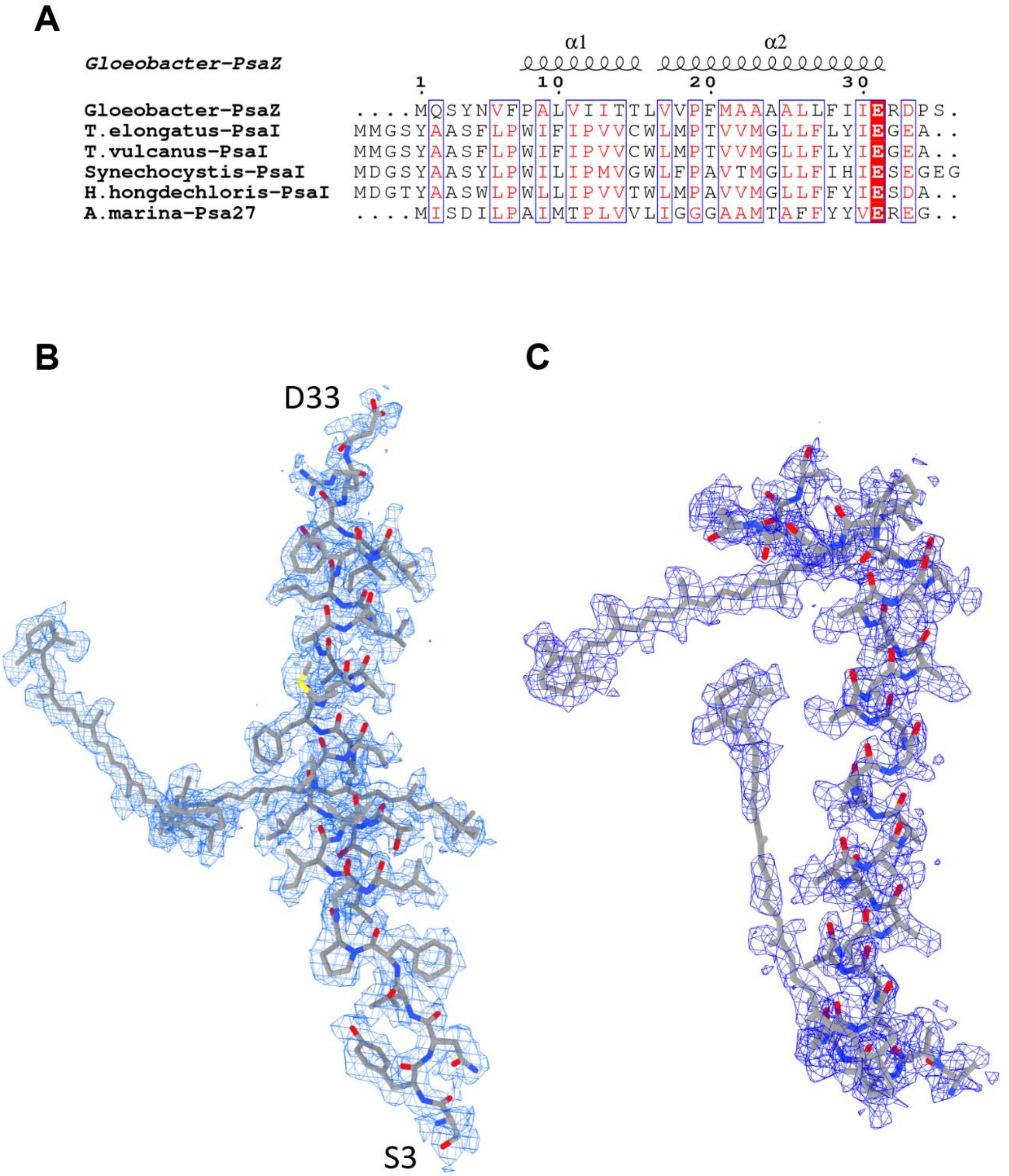


**Fig. S3 Identification of the *Gloeobacter* PSI subunits.** (**A**) Multiple sequence alignment (ClustalW and ESPript) of the *Gloeobacter* PsaZ and other cyanobacterial PsaI subunits. The species shown are *Gloeobacter violaceus* PCC 7421, *Thermosynechococcus elongatus* BP-1, *Thermosynechococcus vulcanus* NIES-2134, *Synechocystis* sp. PCC 6803, *Halomicronema hongdechloris* C2206, *Acaryochloris marina*. (**B**) The density for PsaZ and its corresponding model are shown as blue meshes and gray stick, respectively. (**C**) The density for Unknown subunit and its corresponding model are shown as blue meshes and gray stick, respectively.

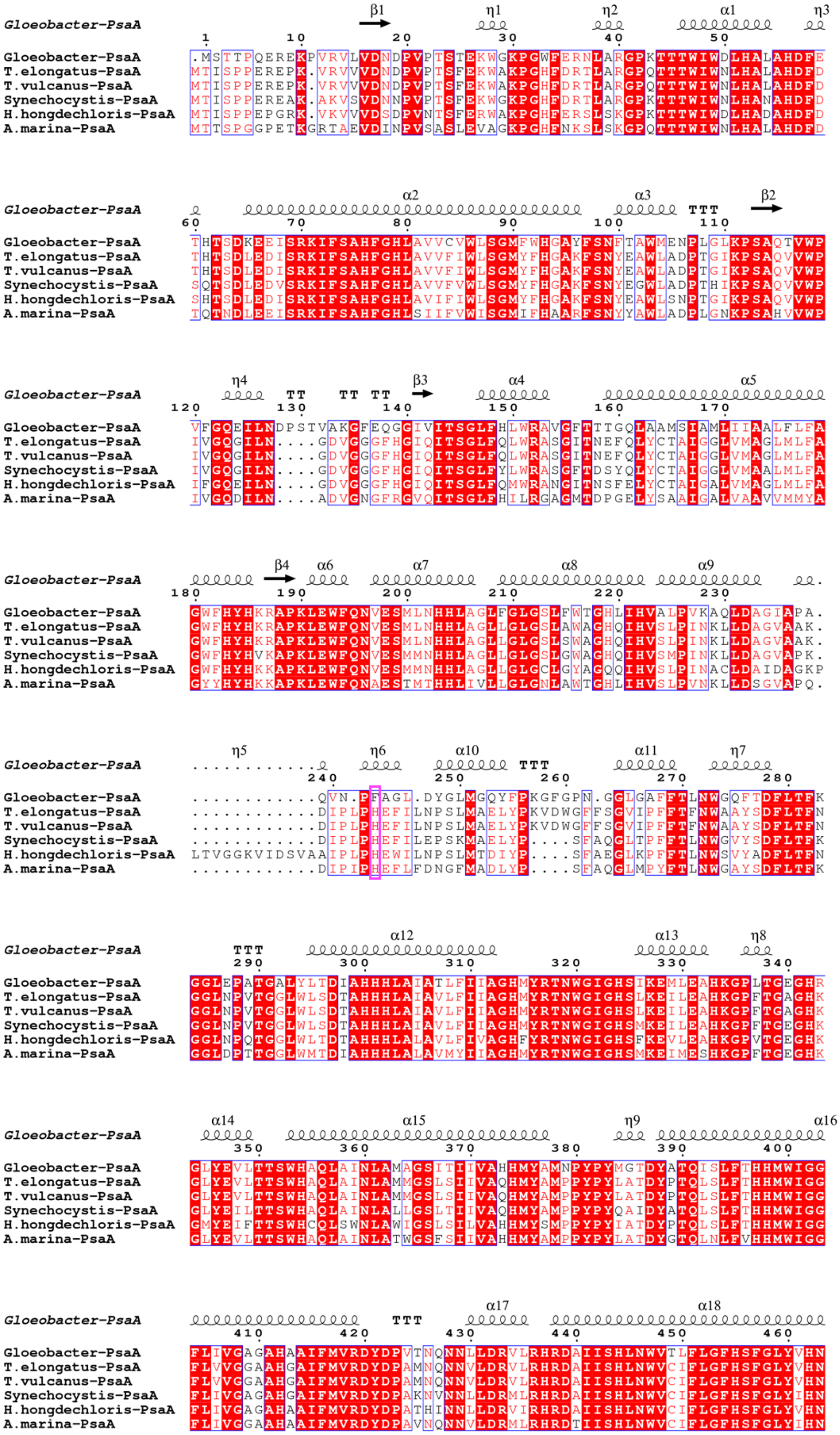
Gloeobacter-PsaA
T.elongatus-PsaA
T.vulcanus-PsaA
Synechocystis-PsaA
H.hongdechloris-PsaA
A.marina-PsaA

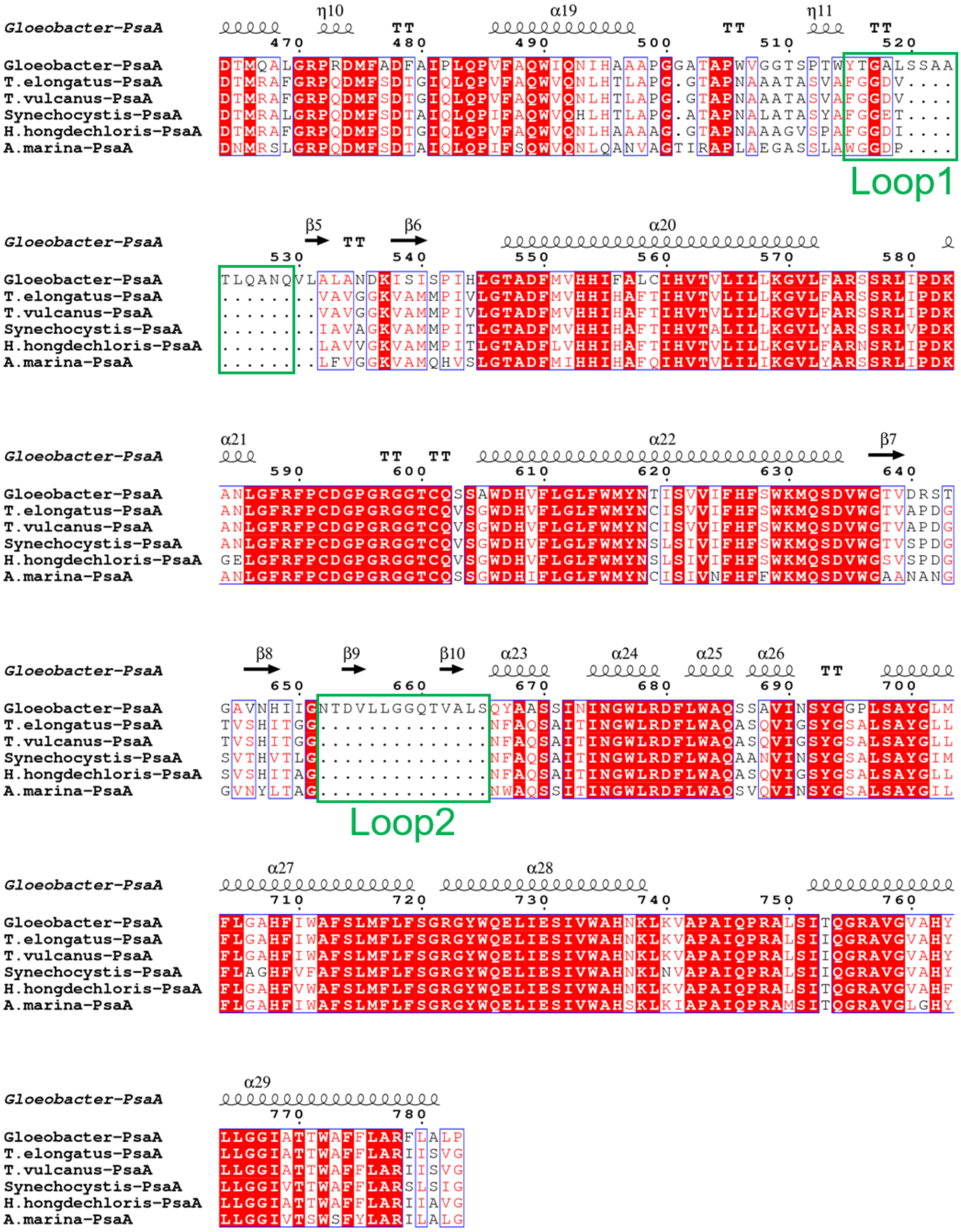

**Fig. S4 Comparison of the structural-known PsaA subunit among 6 species of cyanobacteria.** Multiple sequence alignment was carried out using ClustalW and ESPript. Secondary-structural elements are shown above the sequence. Completely conserved residues are highlighted in red. Loop1 (Tyr515-Gln529) and Loop2 (Asn652-Ser665) are labeled and indicated with green boxes. The pink box stands for the histidine residue involved in the binding of Chl1A. The species shown are *Gloeobacter violaceus* PCC 7421, *Thermosynechococcus elongatus* BP-1, *Thermosynechococcus vulcanus* NIES-2134, *Synechocystis* sp. PCC 6803, *Halomicronema hongdechloris* C2206, *Acaryochloris marina*.

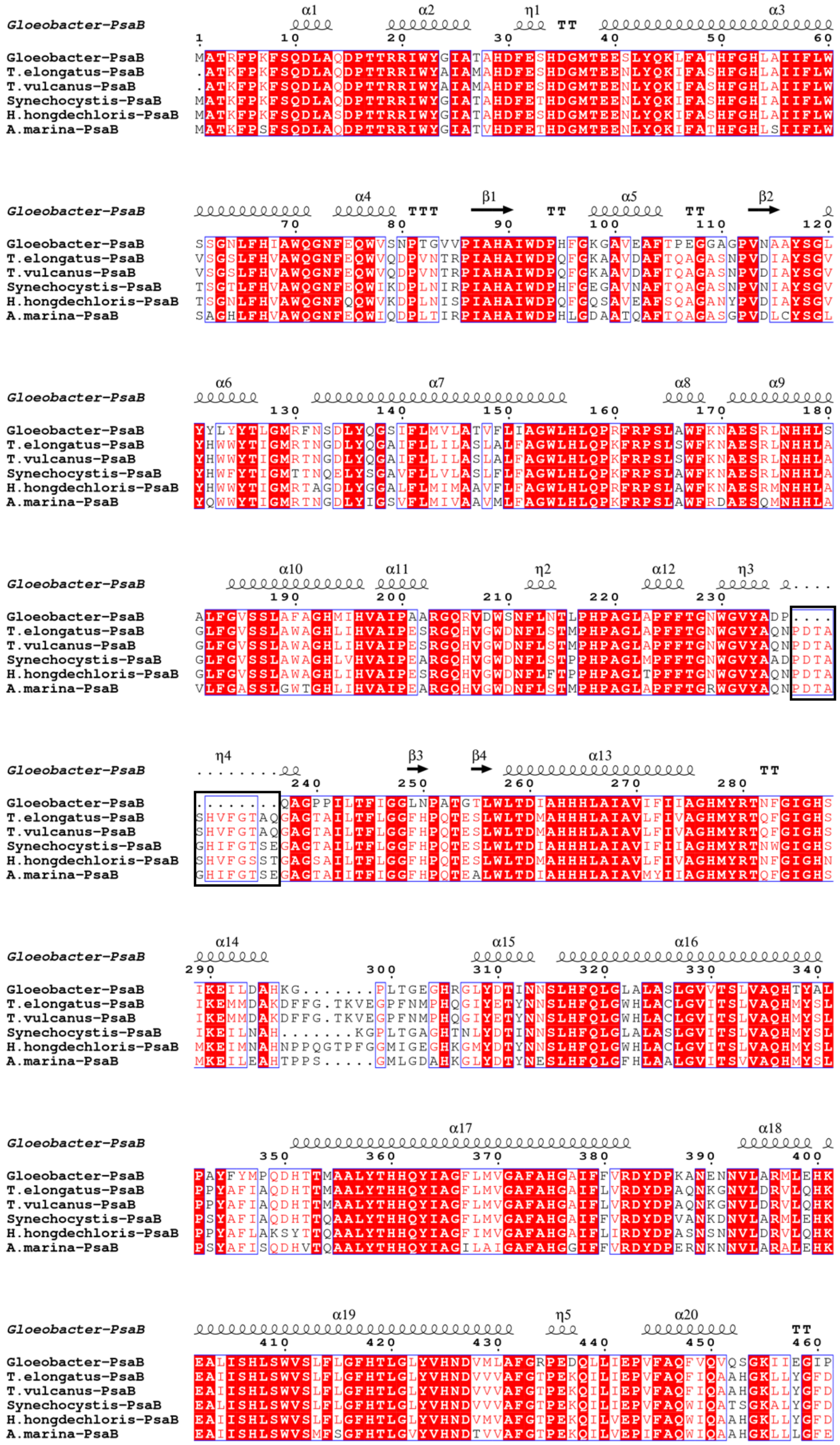
Gloeobacter-PsaB
T.elongatus-PsaB
T.vulcanus-PsaB
Synechocystis-PsaB
H.hongdechloris-PsaB
A.marina-PsaB

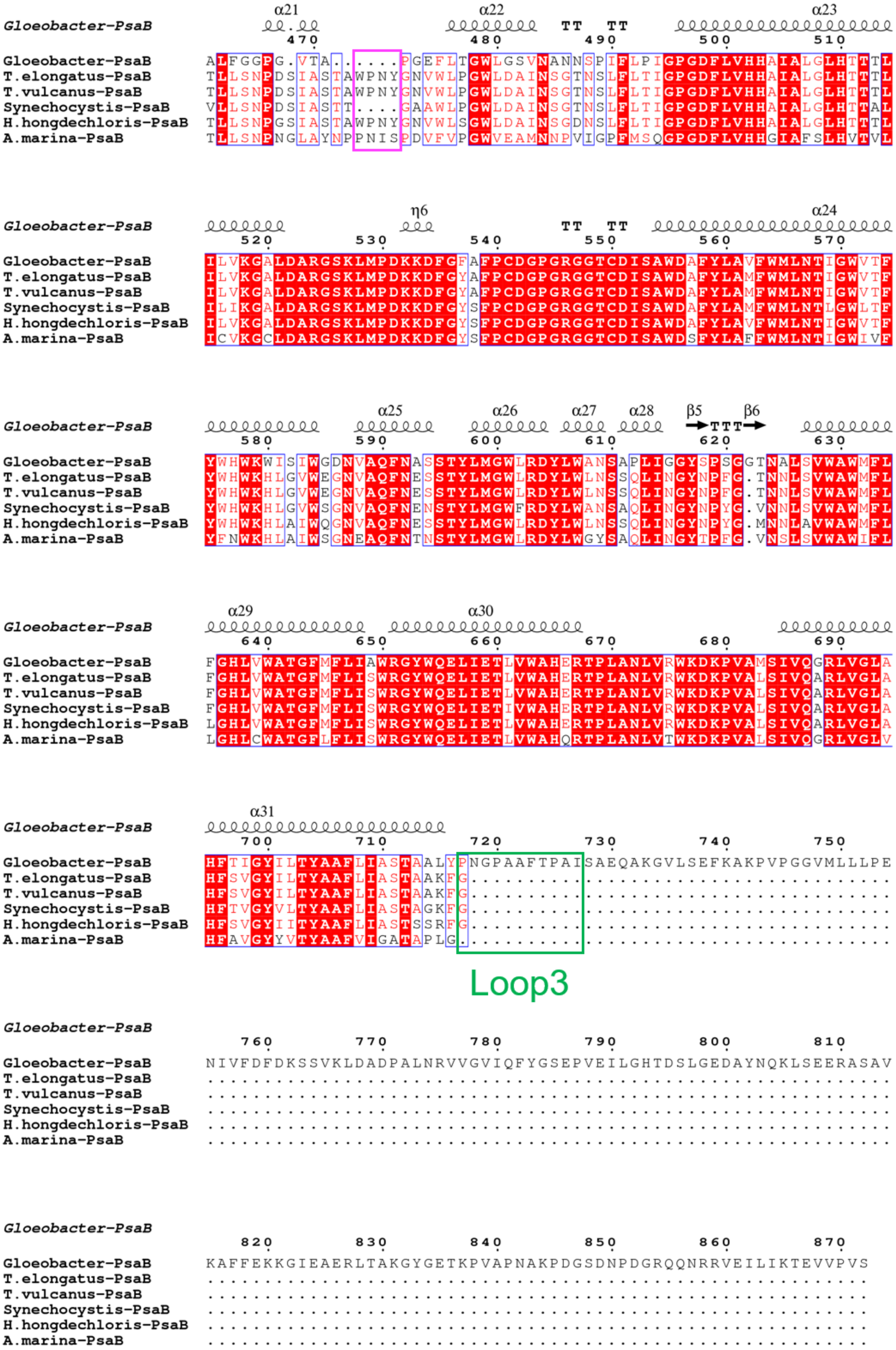
Gloeobacter-PsaB
α21 α22 TT TT α23
470 480 490 500 510
Gloeobacter-PsaB ALFGGPG.VTA......PGEFLTGWLGSVNANNSPIFLPIGPGDFLVHHAIALGLHTTTL
T.elongatus-PsaB TLLSNPDSIASTAWPNYGNVWLPGWLDAINSGTNSLFLTIGPGDFLVHHAIALGLHTTTL
T.vulcanus-PsaB TLLSNPDSIASTAWPNYGNVWLPGWLDAINSGTNSLFLTIGPGDFLVHHAIALGLHTTTL
Synechocystis-PsaB VLLSNPDSIASTT....GAAWLPGWLDAINSGTNSLFLTIGPGDFLVHHAIALGLHTTAL
H.hongdechloris-PsaB TLLSNPGSIASTAWPNYGNVWLSGWLDAINSGDNSLFLTIGPGDFLVHHAIALGLHTTTL
A.marina-PsaB TLLSNPNGLAYNPPNISPDVFVPGWVEAMNNPVIGPFMSQGPGDFLVHHGIAFSLHVTVL
Gloeobacter-PsaB
η6 TT TT α24
520 530 540 550 560 570
Gloeobacter-PsaB ILVKGALDARGSKLMPDKKDFGFAFPCDGPGRGGTCDISAWDAFYLAVFWMLNTIGWVTF
T.elongatus-PsaB ILVKGALDARGSKLMPDKKDFGYAFPCDGPGRGGTCDISAWDAFYLAMFWMLNTIGWVTF
T.vulcanus-PsaB ILVKGALDARGSKLMPDKKDFGYAFPCDGPGRGGTCDISAWDAFYLAMFWMLNTIGWVTF
Synechocystis-PsaB ILIKGALDARGSKLMPDKKDFGYSFPCDGPGRGGTCDISAWDAFYLAMFWMLNTLGWLTF
H.hongdechloris-PsaB ILVKGALDARGSKLMPDKKDFGYSFPCDGPGRGGTCDISAWDAFYLAVFWMLNTIGWVTF
A.marina-PsaB ICVKGCLDARGSKLMPDKKDFGYSFPCDGPGRGGTCDISAWDSFYLAFFWMLNTIGWIVF
Gloeobacter-PsaB
α25 α26 α27 α28 β5 β6
TTT
580 590 600 610 620 630
Gloeobacter-PsaB YWHWKWISIWGDNVAQFNASSTYLMGWLRDYLWANSAPLIGGYSPSGGTNALSVWAWMFL
T.elongatus-PsaB YWHWKHLGVWEGNVAQFNESSTYLMGWLRDYLWLNSSQLINGYNPFG.TNNLSVWAWMFL
T.vulcanus-PsaB YWHWKHLGVWEGNVAQFNESSTYLMGWLRDYLWLNSSQLINGYNPFG.TNNLSVWAWMFL
Synechocystis-PsaB YWHWKHLGVWSGNVAQFNENSTYLMGWFRDYLWANSAQLINGYNPYG.VNNLSVWAWMFL
H.hongdechloris-PsaB YWHWKHLAIWQGNVAQFNESSTYLMGWLRDYLWLNSSQLINGYNPYG.MNNLAVWAWMFL
A.marina-PsaB YFNWKHLAIWSGNEAQFNTNSTYLMGWLRDYLWGYSAQLINGYTPFG.VNSLSVWAWIFL
Gloeobacter-PsaB
α29 α30
640 650 660 670 680 690
Gloeobacter-PsaB FGHLVWATGFMFLIAWRGYWQELIETLVWAHERTPLANLVRWKDKPVAMSIVQGRLVGLA
T.elongatus-PsaB FGHLVWATGFMFLISWRGYWQELIETLVWAHERTPLANLVRWKDKPVALSIVQARLVGLA
T.vulcanus-PsaB FGHLVWATGFMFLISWRGYWQELIETLVWAHERTPLANLVRWKDKPVALSIVQARLVGLA
Synechocystis-PsaB FGHLVWATGFMFLISWRGYWQELIETIVWAHERTPLANLVRWKDKPVALSIVQARLVGLA
H.hongdechloris-PsaB LGHLVWATGFMFLISWRGYWQELIETLVWAHERTPLANLVRWKDKPVALSIVQARLVGLA
A.marina-PsaB LGHLCWATGFLFLISWRGYWQELIETLVWAHQRTPLANLVTWKDKPVALSIVQGRLVGLV
Gloeobacter-PsaB
α31
700 710 720 730 740 750
Gloeobacter-PsaB HFTIGYILTYAAFLIASTAALYPNGPAAFTPAISAEQAKGVLSEFKAKPVPGGVMLLLPE
T.elongatus-PsaB HFSVGYILTYAAFLIASTAAKFG.....................................
T.vulcanus-PsaB HFSVGYILTYAAFLIASTAAKFG.....................................
Synechocystis-PsaB HFTVGYVLTYAAFLIASTAGKFG.....................................
H.hongdechloris-PsaB HFSVGYIITYAAFLIASTSSRFG.....................................
A.marina-PsaB HFAVGYYVTYAAFVIGATAPLG......................................
Loop3
Gloeobacter-PsaB
760 770 780 790 800 810
Gloeobacter-PsaB NIVFDFDKSSVKLDADPALNRVVGVIQFYGSEPVEILGHTDSLGEDAYNQKLSEERASAV
T.elongatus-PsaB ............................................................
T.vulcanus-PsaB ............................................................
Synechocystis-PsaB ............................................................
H.hongdechloris-PsaB ............................................................
A.marina-PsaB ............................................................
Gloeobacter-PsaB
820 830 840 850 860 870
Gloeobacter-PsaB KAFFEKKGIEAERLTAKGYGETKPVAPNAKPDGSDNPDGRQQNRRVEILIKTEVVPVS
T.elongatus-PsaB ..........................................................
T.vulcanus-PsaB ..........................................................
Synechocystis-PsaB ..........................................................
H.hongdechloris-PsaB ..........................................................
A.marina-PsaB ..........................................................

**Fig. S5 Comparison of the structural-known PsaB subunit among 6 species of cyanobacteria.** Multiple sequence alignment was carried out using ClustalW and ESPript. Secondary-structural elements are shown above the sequence. Secondary-structural elements are shown above the sequence. Completely conserved residues are highlighted in red. Loop3 (Pro717-Ile727) is labeled and indicated with green box. The pink box indicates the loop involved in the binding of Chl1B. The black box stands for the deletion in the *Gloeobacter* PsaB. The species shown are *Gloeobacter violaceus* PCC 7421, *Thermosynechococcus elongatus* BP-1, *Thermosynechococcus vulcanus* NIES-2134, *Synechocystis* sp. PCC 6803, *Halomicronema hongdechloris* C2206, *Acaryochloris marina*.

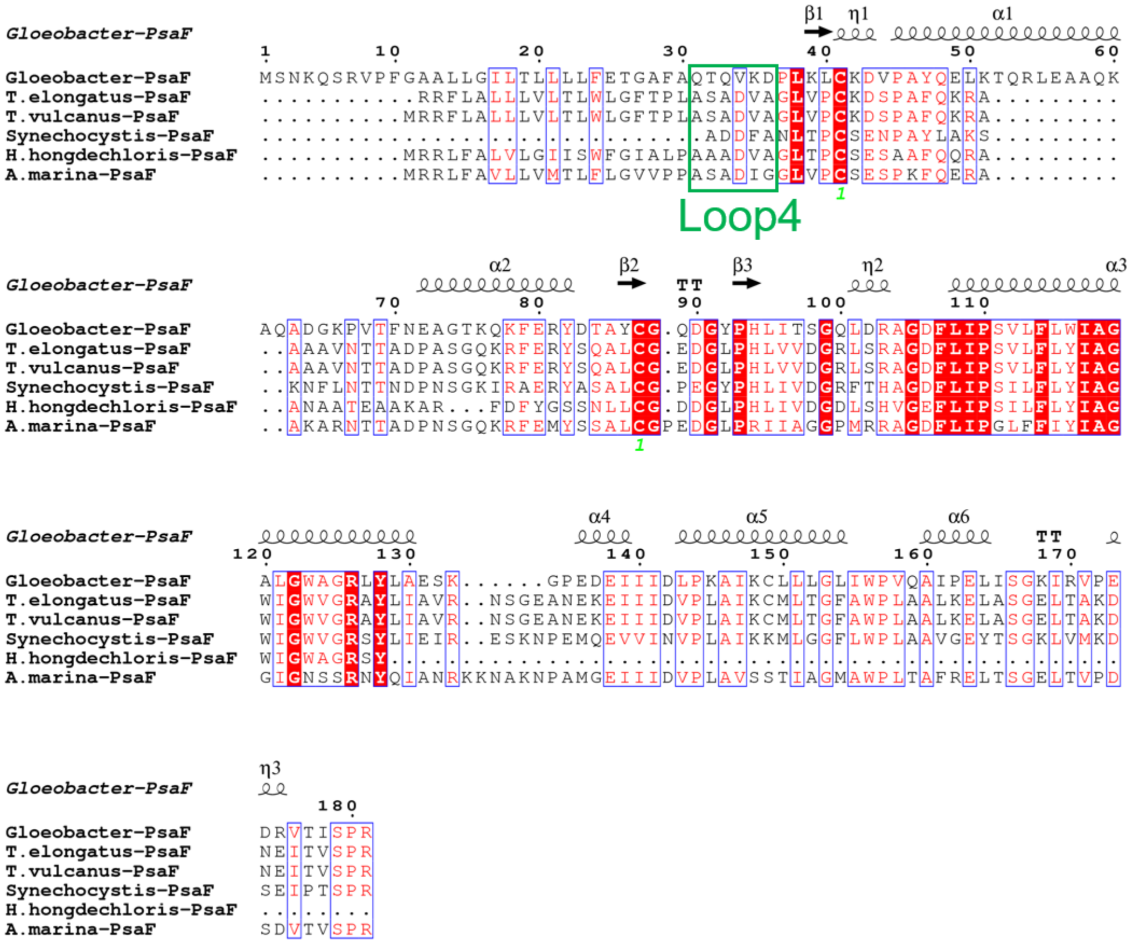

**Fig. S6 Comparison of the structural-known PsaF subunit among 6 species of cyanobacteria.** Multiple sequence alignment was carried out using ClustalW and ESPript. Secondary-structural elements are shown above the sequence. Completely conserved residues are highlighted in red. Loop4 (Gln31-Asp36) is labeled and indicated with green box. The species shown are *Gloeobacter violaceus* PCC 7421, *Thermosynechococcus elongatus* BP-1, *Thermosynechococcus vulcanus* NIES-2134, *Synechocystis* sp. PCC 6803, *Halomicronema hongdechloris* C2206, *Acaryochloris marina*.

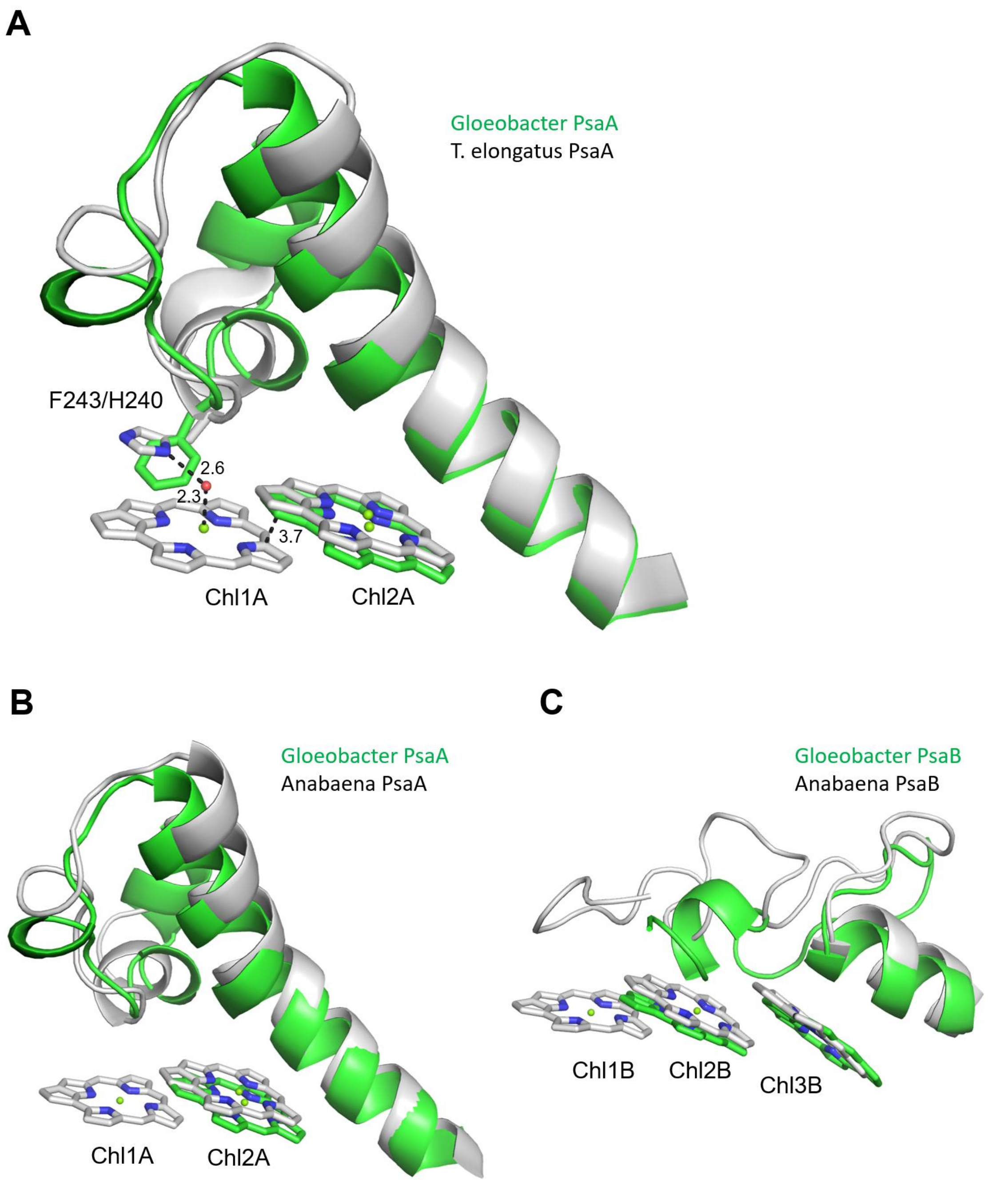


**Fig. S7 Structural comparison of Low1 and Low2 site in PsaA between *Gloeobacter* and other organisms.** (**A**) Superposition of Low1 site in the *Gloeobacter* PSI (green) with the *T. elongatus* PSI (gray). Phe243 of the *Gloeobacter* PsaA and His240 of the *T. elongatus* PsaA are shown in stick model. Interactions are indicated by dashed lines with distances labeled in Å. (**B**, **C**) Superposition of the Low1 site (**B**) and Low2 site (**C**) in the *Gloeobacter* PSI (green) with the *Anabaena* PSI (gray).

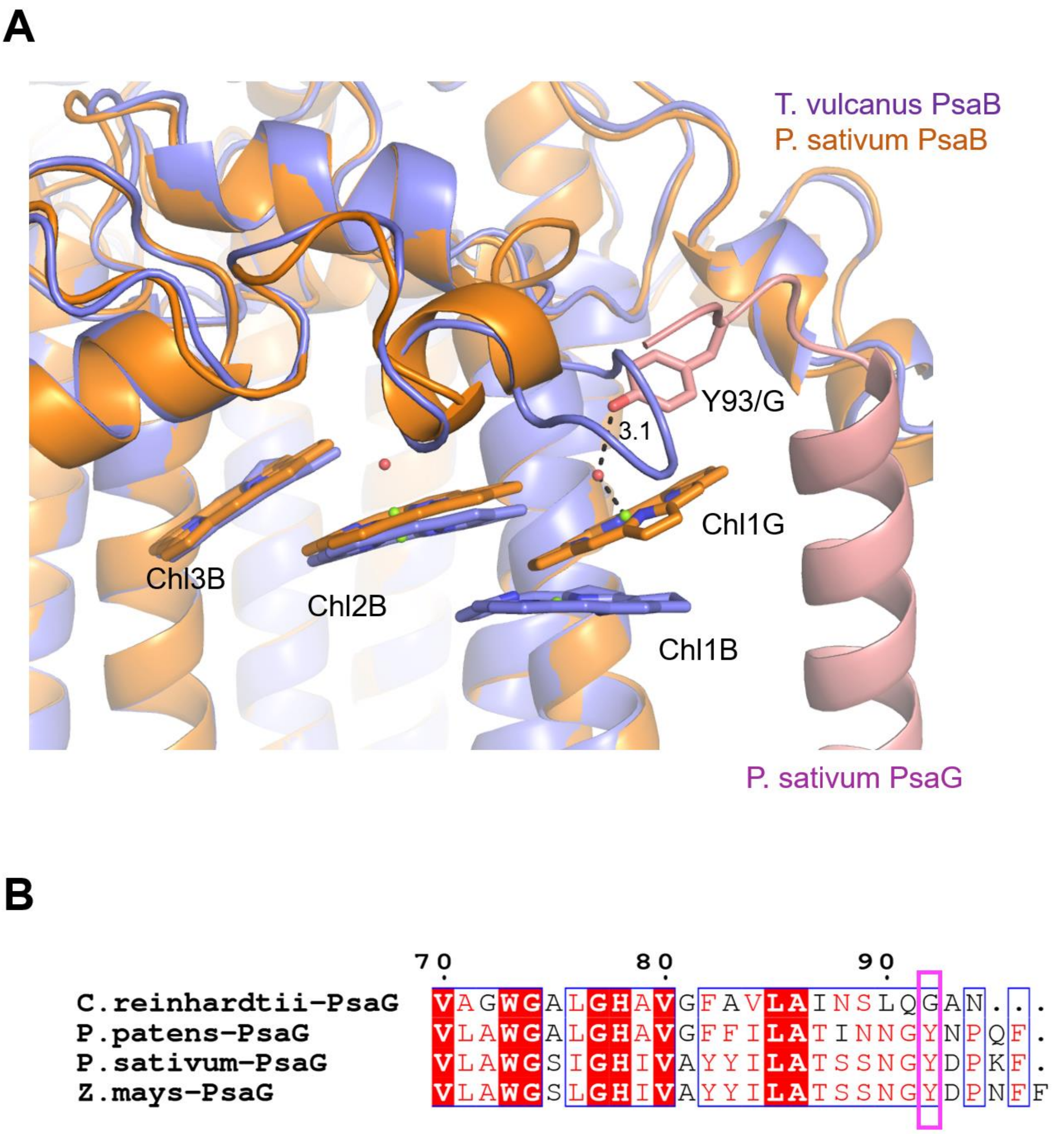


**Fig. S8 Structural comparisons of the Low2 site.** (**A**) Superposition of the Low2 site in the *T. vulcanus* PsaB (purple) with the *P. sativum* PsaB (orange). The *P. sativum* PsaG is colored pink. Tyr93 involved in the binding of Chl1G is shown in stick model. (**B**) Multiple sequence alignment of PsaG using ClustalW and ESPript. The species shown are *Chlamydomonas reinhardtii*, *Physcomitrella patens*, *Pisum sativum*, and *Zea mays*. The pink box displays the tyrosine residue involved in the binding of Chl1G.

**Table S1. Cryo-EM data collection and structural analysis statistics.**

| | |
|---|---|
| Complex | PSI trimer |
| PDB ID | 7F4V |
| EMDB ID | EMD-31455 |
| Data collection and processing | |
| Magnification | 60000 |
| Voltage (kV) | 300 |
| Electron exposure ($e^{-}$/Å) | 70.22 |
| Defocus range (μm) | −1.8 to −0.6 |
| Pixel size (Å) | 0.823 |
| Symmetry imposed | C3 |
| Initial particle number | 961960 |
| Final particle number | 261743 |
| Map resolution (Å) | 2.04 |
| FSC threshold | 0.143 |
| Refinement | |
| Initial Model used (PDB code) | Homology modeling |
| Model resolution (Å) | 2.01 |
| FSC threshold | 0.5 |
| Map sharpening B factor ($Å^2$) | −58.0 |
| Model composition (trimer) | |
| Non-hydrogen atoms | 67641 |
| Protein | 49899 |
| Ligand | 17340 |
| Water | 402 |
| B factors ($Å^2$) | |
| Protein | 39.2 |
| Ligand | 40.5 |
| Water | 27.3 |
| R.m.s deviations | |
| Bond lengths (Å) | 0.025 |
| Bond angles (°) | 2.45 |
| Validation | |
| MolProbity score | 1.78 |
| Clashscore | 4.77 |
| Poor rotamers (%) | 2.78 |
| EMRinger score | 6.48 |
| Ramachandran plot | |

| | |
|---|---|
| Favored (%) | 96.80 |
| Allowed (%) | 3.11 |
| Disallowed (%) | 0.09 |

**Table S2. Averaged Q-score in each subunit and cofactors in each monomer unit of the PSI core.**

| Protein | Amino acid residues | Averaged Q-score | Chlorophyll | Carotenoid | Lipid | Others |
|---|---|---|---|---|---|---|
| **PsaA** | 11–782 | 0.85 | 43 Chl *a*<br>1 Chl *a′* | 5 BCR | 2 LHG | 1 [4Fe-4S] cluster,<br>1 menaquinone-4 |
| **PsaB** | 3–727 | 0.87 | 41 Chl *a* | 7 BCR | 1 LMG<br>1 LHG | 1 menaquinone-4 |
| **PsaC** | 2–81 | 0.87 | | | | 2 [4Fe-4S] cluster |
| **PsaD** | 11–143 | 0.81 | | | | |
| **PsaE** | 3–64 | 0.76 | | | | |
| **PsaF** | 31–181 | 0.76 | 1 Chl *a* | 1 BCR | | |
| **PsaZ** | 3–33 | 0.85 | | 2 BCR | | |
| **Unknown** | 1–33 | 0.80 | | 2 BCR | | |
| **PsaL** | 11–782 | 0.85 | 3 Chl *a* | 2 BCR | | |
| **PsaM** | 3–727 | 0.84 | | 1 BCR | | |
| **Total** | | | 89 | 20 | 4 | 5 |

BCR, *β*-carotene; LMG, distearoylmonogalactosyl diglyceride; LHG, dipalmitoylphosphatidyl glycerol.
